# Cognitive self-regulation influences pain-related physiology

**DOI:** 10.1101/361519

**Authors:** Gordon Matthewson, Choong-Wan Woo, Marianne C. Reddan, Tor D. Wager

**Affiliations:** Department of Psychology and Neuroscience, University of Colorado, Boulder, USA; Institute of Cognitive Science, University of Colorado, Boulder, USA; Center for Neuroscience Imaging Research, Institute for Basic Science, Suwon, South Korea; Department of Biomedical Engineering, Sungkyunkwan University, Suwon, South Korea

**Keywords:** pain, self-regulation, autonomic nervous system, SCR, ECG

## Abstract

Cognitive self-regulation can shape pain experience, but its effects on autonomic responses to painful events are unclear. In this study, participants (*N* = 41) deployed a cognitive strategy based on reappraisal and imagination to regulate pain up or down on different trials while skin conductance responses (SCR) and electrocardiogram (ECG) activity were recorded. Using a machine learning approach, we first developed stimulus-locked SCR and ECG physiological markers predictive of pain ratings. The physiological markers demonstrated high sensitivity and moderate specificity in predicting pain across two datasets, including an independent test dataset (*N* = 84). When we tested the markers on the cognitive self-regulation data, we found that cognitive self-regulation had significant impacts on both pain ratings and pain-related physiology in accordance with regulatory goals. These findings suggest that self-regulation can impact autonomic nervous system responses to painful stimuli and provide pain-related autonomic profiles for future studies.

## Introduction

Cognitive self-regulation is a way of modulating pain and emotion by consciously changing one’s thoughts and appraisals of sensations and the context in which they occur [1; 10; 19; 26; 27; 36]. Psychological interventions such as hypnosis and placebo have long been documented as effective methods of pain control [31], and several cognitive self-regulation techniques have also been documented for their ability to reduce pain (for a review, see [15]). Some of the most prominent include mental imagery [6; 11] and *reappraisal*, which involves contextual reinterpretation of painful sensations [39; 42]. Beliefs and conditioning are known to have strong physiological impacts, such as in the case of placebo effects [24; 33; 47], but the relationship between conscious self-regulation and autonomic responses remains less understood. Here, we studied whether conscious, top-down self-regulation can impact pain-related autonomic physiology.

Painful events induce dramatic changes in the autonomic nervous system. These changes, including increases in blood pressure, heart rate, skin conductance, and pupil dilation [4; 8; 9; 18; 34], are consistent with sympathetic activation and parasympathetic withdrawal and thought to be mediated by interactions with parabrachial nociceptive pathways in the brainstem [5; 7; 40]. However, quantifying pain-related autonomic responses in the context of cognitive pain modulation is challenging because autonomic changes are not specific to pain. During cognitive pain modulation, for example, the autonomic nervous system responds to noxious stimulation, but also to orientation to a stimulus [16], cognitive load [32; 35], and stress [22]. As a result, it is difficult to isolate cognitive effects on pain-related physiology from those related to other processes, including cognitive regulation itself [44; 45]. For example, regulation-induced reductions in pain-related autonomic responses could be masked by increases due to the cognitive demands of regulation itself, resulting in a null net effect [13]. Therefore, to quantify the effects of cognitive regulation on pain physiology, there is a need to first identify components of autonomic responses that are as tightly linked to pain as possible, and then test the effects of regulation on these identified component measures. In EEG research, for example, component-based processes (e.g., Independent Components Analysis) are routinely used to decompose EEG responses into separate components, some of which reflect artifactual signals and others which reflect multiple task-related signals of interest [30]. To our knowledge, however, this approach has not been applied to autonomic responses.

In the current study, we aimed to examine whether self-regulation influences pain-related physiology by developing pain-predictive physiological markers based on skin conductance response (SCR) and electrocardiogram (ECG) data. We reasoned that if pain-related autonomic signals could be isolated by extracting a temporal waveform (component) optimized to predict pain, it could provide a better test of whether cognitive regulation reduces this autonomic signal (see Fig. 1a). In **Study 1**, 41 participants engaged in self-regulation to increase or decrease pain while experiencing six different levels of painful heat (44.3–49.3 °C in 1°C increments). Using data only from trials in which participants passively experienced thermal pain with no regulation instructions, we developed stimulus-locked SCR and ECG models predictive of pain ratings using principal component regression [20] (Analysis 1 in Fig. 1b). This first phase was designed to minimize influences of psychological processes (e.g., expectations and self-regulation). In **Study 2**, the resulting pain-related SCR measure (the strongest pain-predictive signal) were validated on an independent study dataset with 42 pairs of romantic partners (total *N* = 84; Analysis 2 in Fig. 1b). Here, three levels of painful heat (47, 48, and 49 °C) were delivered to one participant in each pair (pain receiver), and the other person observed his or her partner experiencing pain (pain observer). SCRs were simultaneously recorded in both participants throughout. This design allowed us to assess the SCR measure’s provisional sensitivity (response to first-person experience of pain) and specificity (response to observed pain, which is a non-painful, but salient event) in an independent dataset [43]. Lastly, we applied the physiological pain markers to data during cognitive self-regulation in **Study 1** to test whether cognitive self-regulation affects SCR and ECG-based pain-predictive physiological measures (Analysis 3 in Fig. 1b). We found that whereas cognitive regulation had no effect on autonomic responses in traditional analyses of event-related averages, it exerted bidirectional influences on autonomic measures optimized to be pain-predictive.

**Figure 1.**
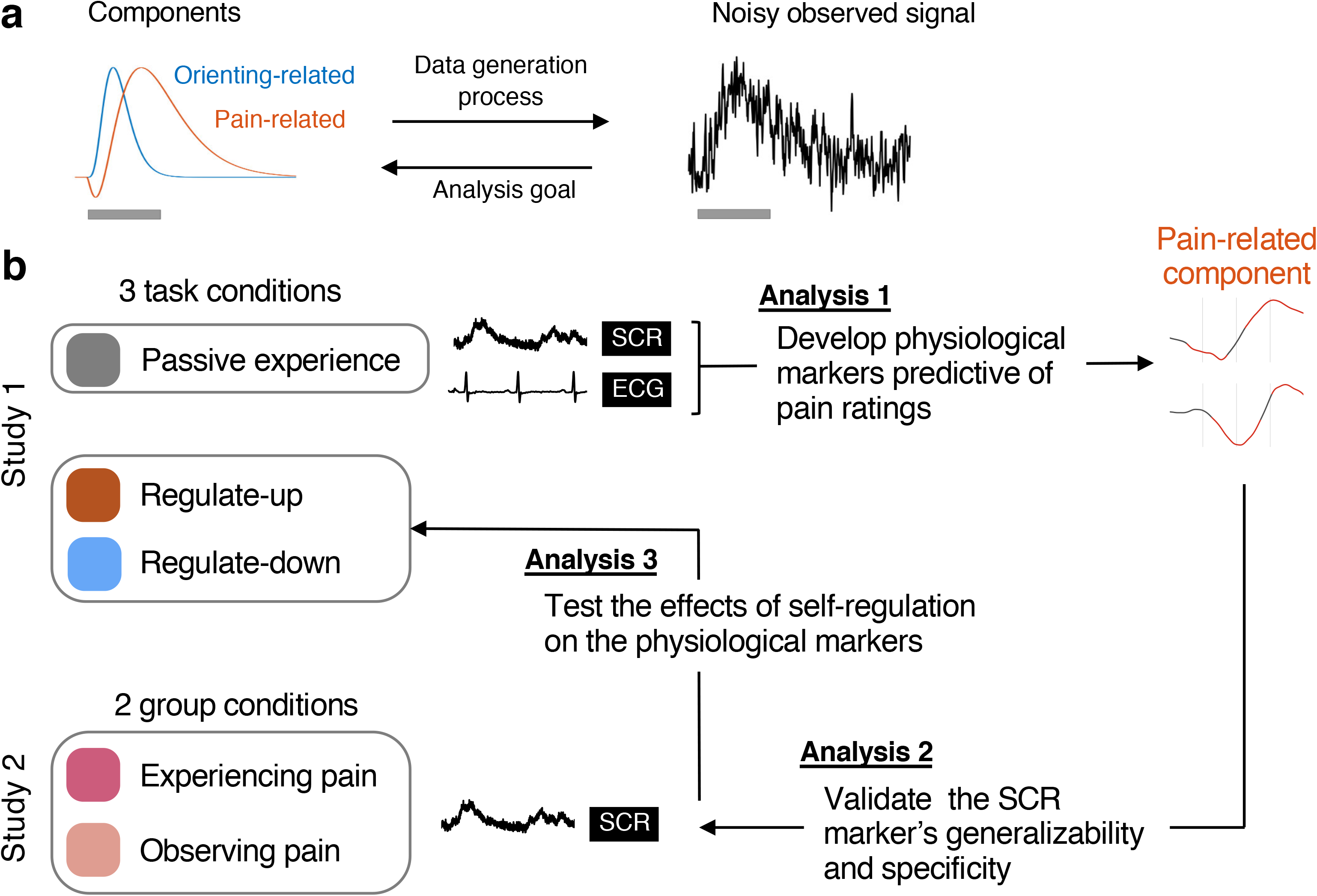
Concept and analysis pipeline. **(a)** Concept behind component-based analysis. Like other complex signals, autonomic responses may be composed of multiple underlying signals mixed together. Here, two components are illustrated, one unrelated and another related to pain reports. The observed signal is a mixture of the two plus noise. The gray bar shows the stimulus period. The goal of component-based analysis is to recover the underlying signals. In this case, we aimed to identify a waveform correlated with pain reports, separating this to the extent possible from non-pain-related signal and noise components. This allowed us to test self-regulation effects on the amplitude of the pain-related component for each participant. **(b)** The analysis plan includes three steps. **Analysis 1:** We developed pain-predictive measures based on SCR and ECG using Study 1 (*N* = 41) data from the passive experience condition (no regulation). We used leave-one-subject-out cross-validation to estimate their accuracy in predicting pain when applied to new participants. **Analysis 2:** We validated the SCR marker with an independent dataset (Study 2, *N* = 84) to establish its provisional generalizability and specificity. We tested the marker on participants experiencing pain and observing their romantic partner experience pain. **Analysis 3:** We applied the pain-related autonomic measures to data collected during cognitive self-regulation in Study 1, to test whether self-regulation changed pain-predictive physiological responses. We used cross-validation, so that the measures were only applied to subjects not used in measure development.

## Methods

### Participants

#### Study 1

42 healthy participants with no history of psychiatric, neurological, or pain disorders and no current pain were recruited for this experiment. A sample size of 42 was chosen to both ensure sufficient statistical power and minimize the order effects due to the different condition types (for the randomization procedure, see Task Design). Based on the effect size estimates from the previous study [52] (Cohen’s *d* = 0.70 for the self-regulation effect on self-reported pain), a sample size of 42 was estimated to provide 99.2% power.

Participants were recruited through Craigslist.org and advertisements placed on the University of Colorado campus, and further contacted through telephone and email. One participant decided to stop the experiment halfway through because his skin was becoming too sensitive, leaving a final sample size of *N*=41 (20 females, 21 males; age = 24.3 ± 5.6 [mean ± SD] years; range: 18-41 years). 36 participants were of Caucasian ethnicity, 2 participants Hispanic, 1 African-American, 1 Asian, and 1 participant reported being mixed ethnicity. All participants gave written informed consent and were compensated $12 an hour for their participation.

#### Study 2

48 romantic couples (N = 96) with no history of psychiatric, neurological, or pain disorders and no current pain participated together in this experiment. Six participants from different couples had technical issues in SCR signal acquisition, leaving a final sample size of 42 couples (*N* = 84) as either the main participant of *N* = 42 (21 females, age = 27.90 ± 6.29 years, range = 21 − 47), who experienced pain, or the partner *N* = 42 (22 females, age = 27.45 ± 6.20 years, range = 21 − 47), who did not experience pain, but observed their partners experiencing pain. 38 participants were of Caucasian ethnicity, 7 Hispanic, 1 African American, 3 Native American, and 2 Asian American (and 33 preferred not to respond). All participants gave written informed consent and were compensated $12 an hour for their participation.

### Thermal stimulation

Thermal stimulation was delivered to participants using an ATS Pathway System (Medoc Ltd.) with a 16-mm Peltier thermode end-plate.

#### Study 1

Heat stimulations were delivered to three sites located on the middle forearm that alternated between runs. Each stimulation lasted 12.5 seconds, with 3-second ramp-up and 2-second ramp-down periods and 7.5 seconds at target temperature. Six levels of temperature were administered to the participants (level 1: 44.3°C; level 2: 45.3°C; level 3: 46.3°C; level 4: 47.3°C; level 5: 48.3°C; level 6: 49.3°C).

#### Study 2

Heat stimulations were delivered to three sites located on the participants’ left leg. Each stimulation lasted 12 seconds, with 3.5-second ramp-up and 1-second ramp-down periods and 7.5 seconds at target temperature. Three levels of temperature were administered to the participants (level 1: 47°C; level 2: 48°C; level 3: 49°C).

### Rating scales

In Study 1 and 2, we used the same general Labeled Magnitude Scale (gLMS) to assess pain intensity and unpleasantness [3]. We used gLMS because it provides more valid across-group comparisons and more effectively captures variance in the high-pain range than the visual analog or categorical scales. In the pain intensity gLMS the anchors began with “No sensation” (0) to the far left of the scale, and continued to the right in a graded fashion with anchors of “Barely detectable” (1.4), “Weak” (6.1), “Moderate” (17.2), “Strong” (35.4), and “Very strong” (53.3), until “Strongest imaginable sensation of any kind” (100) on the far right. Whereas the pain intensity scale progressed in a unidirectional fashion from left to right, the pain unpleasantness scale was used in a bidirectional fashion, with “Neutral” in the center, increasing unpleasantness progressing to the left, and increasing pleasantness progressing to the right. The same increments from the first scale were used in each direction, with the end anchor “Strongest unpleasantness imaginable of any kind” to the left, and “Strongest pleasantness imaginable of any kind” to the far right. The length of the scales was proportional such that the pain intensity scale was exactly half that of the pain unpleasantness scale. During the main task, the intermediate anchors were removed to eliminate anchor effects [21].

### General procedure

#### Study 1

Participants were given a brief overview of the experiment, which explained that they were participating in a study on the physiological effects of cognitive pain regulation. After participants provided informed consent, we explained the gLMS rating scales used throughout the experiment to the participant and allowed them to practice using the scales. After a verbal explanation was given of what each anchor signified, participants were asked to explain the scale back to the experimenter to ensure that the participants understood the scale correctly.

Skin sites were then selected for stimulation based upon a calibration procedure. During this procedure, pain intensity ratings were collected from a 47.3°C and a 48.3° C stimulation to eight different sites on the forearm to determine which sites on the arm produced the most reliable and similar pain ratings, and additionally to ensure that the heat was indeed painful, but not intolerable or excessive. The sites of the stimulations were randomized between eight different locations evenly spaced between the wrist and the elbow on the volar surface of the left forearm. Three sites that the participant rated most similarly were chosen for use in the main procedure.

Following the calibration procedure was a regulation practice session, in which the experimenter asked the participant to relax, close their eyes, and follow along with a script read aloud by the experimenter designed to promote awareness of sensations and cognitive control over one’s sensations (see **Supplemental Methods** for a full practice script). Participants were informed that an effective way to manipulate pain is to change the meaning of painful sensations, and then led through instructions designed to increase or decrease the experience of pain (see **“**Cognitive Regulation Instructions” below). These instructions were designed to give participants confidence in their ability to regulate pain, as this is essential for any self-regulation technique to be effective. A full practice script can be found in the supplemental material.

The main task was grouped as 9 runs of 6 thermal stimulations each. There were three run conditions: an up-regulation condition, a down-regulation condition, and a passive control condition. Regulation condition for each run was pseudorandomized using a Latin-square method, resulting in six different sets of run orders, one of which was assigned to a participant before the experiment began. Each run began with a stimulation of 49.3° C to minimize the effects of within-run sensitization and habituation to heat [23]. After this stimulation, the regulation instructions for the run were shown on screen.

After it was clear that the participant understood the regulation instructions for the run, the stimulations began. The timing of a single trial can be seen in Fig. S1. Six temperatures between 44.3-49.3°C were administered in a randomized order, and after each heat stimulation pain intensity and unpleasantness ratings were collected. The order in which the two rating scales were presented was randomized. After three trials, a reminder screen was presented, which provided encouragement and reminded the participant which type of regulation they were supposed to be using.

#### Study 2

Couples provided informed consent, and then each member of the couple was randomly assigned to be either the main participant, who experienced pain, or the partner, who did not experience pain but provided support. Specifically, partners observed the main participants receiving painful stimulation (“Present” condition) or provided supportive touch (“Hand-holding” or “Gentle stroking” conditions) (see **Supplemental Methods** for a detailed task design for Study 2). The current study uses data only from the “Present” condition. The main participants underwent the same rating scale introduction as Study 1. Skin site selection was fixed *a priori* before the experiment and was the same for each person (on the outer left leg, right below the knee, in the center of the leg, and right above the ankle). Temperatures were also determined *a priori* to be 47°, 48°, and 49°C.

### Cognitive Regulation Instructions (Study 1)

The regulation instructions (see **Supplemental Materials** for full script) combined (a) instructions designed to enhance participants’ pain regulation self-efficacy [2] (e.g., “you can develop a powerful relationship with your sensations”, “it will become much easier to manipulate your experience once you have stronger sensations to work with”) with (b) instructions to engage in specific forms of imagery targeting several aspects of pain, including appraisals of its intensity and harmfulness.

In the down-regulation condition, participants were instructed to engage in imagery and appraisals that minimized danger and enhanced the counterfactual pleasantness of the stimulation, e.g., “Focus on the part of the sensation that is pleasantly warm, like a blanket on a cold day, and the aspects of the heat that are calming, soothing, and relaxing”; “…turn down the dial of your pain sensation.” They also emphasized imagery that promotes acceptance and disengagement from negative affect, e.g., “allow the pain and heat to be carried away, flowing away from your body”; “visualize the powerful warmth flowing and spreading through you as it gives you energy and life.” In the up-regulation condition, participants were instructed to engage in imagery intended to engage negative affect and enhance appraisals of harm, e.g., “Pay attention to the burning, stinging and shooting sensations”; “imagine how unpleasant the pain is”; “You can use your mind to turn up the dial of the pain”; “visualize your skin sizzling, melting, and bubbling as a result of the intense heat.” In the neutral condition, participants were explicitly instructed not to attempt to regulate the pain, and instead to focus on accurately perceiving the sensations, e.g., “Try not to regulate or change your sensation, but instead accurately rate what each sensation was like as you felt it.”

These strategies were chosen because of their effectiveness in published [6; 11; 15; 52] and ongoing work, and are related to several types of strategies described in the literature. For example, for down-regulation of pain, asking participants to “imagine that the thermal stimulations are less painful than they are” is related to what previous literature has described as *pain acknowledging* [46] or *reinterpretation* [11], whereas asking participants to focus on aspects of the heat that are “pleasantly warm, like a blanket on a cold day” is similar to *pleasant imagery* or *dramatized coping*, which focuses on the narrative context or situational meaning surrounding stimulation [15]. For up-regulation of pain, asking participants to “focus on how unpleasant the pain is” is a negative form of *pain acknowledging* [46], whereas “picture your skin being held up against a glowing hot metal or fire” is related to *dramatized coping* [15]. We intentionally made our self-regulation instructions broad enough to include multiple components of self-regulation strategies because the aim of the current study is to examine the overall effects of self-regulation on pain physiology, not to compare the effects of various self-regulation strategies. Also, an important commonality between the up-regulation and down-regulation instructions is their emphasis on consciously attending to the stimuli and changing their meaning, instead of directing attention elsewhere, such as in distraction-based pain regulation strategies.

Participants engaged in a post-experiment description of the strategies they actually used, which were coded in relation to eight common strategies described in the literature. An analysis of the strategies used by participants can be found in Fig. S2.

**Figure 2.**
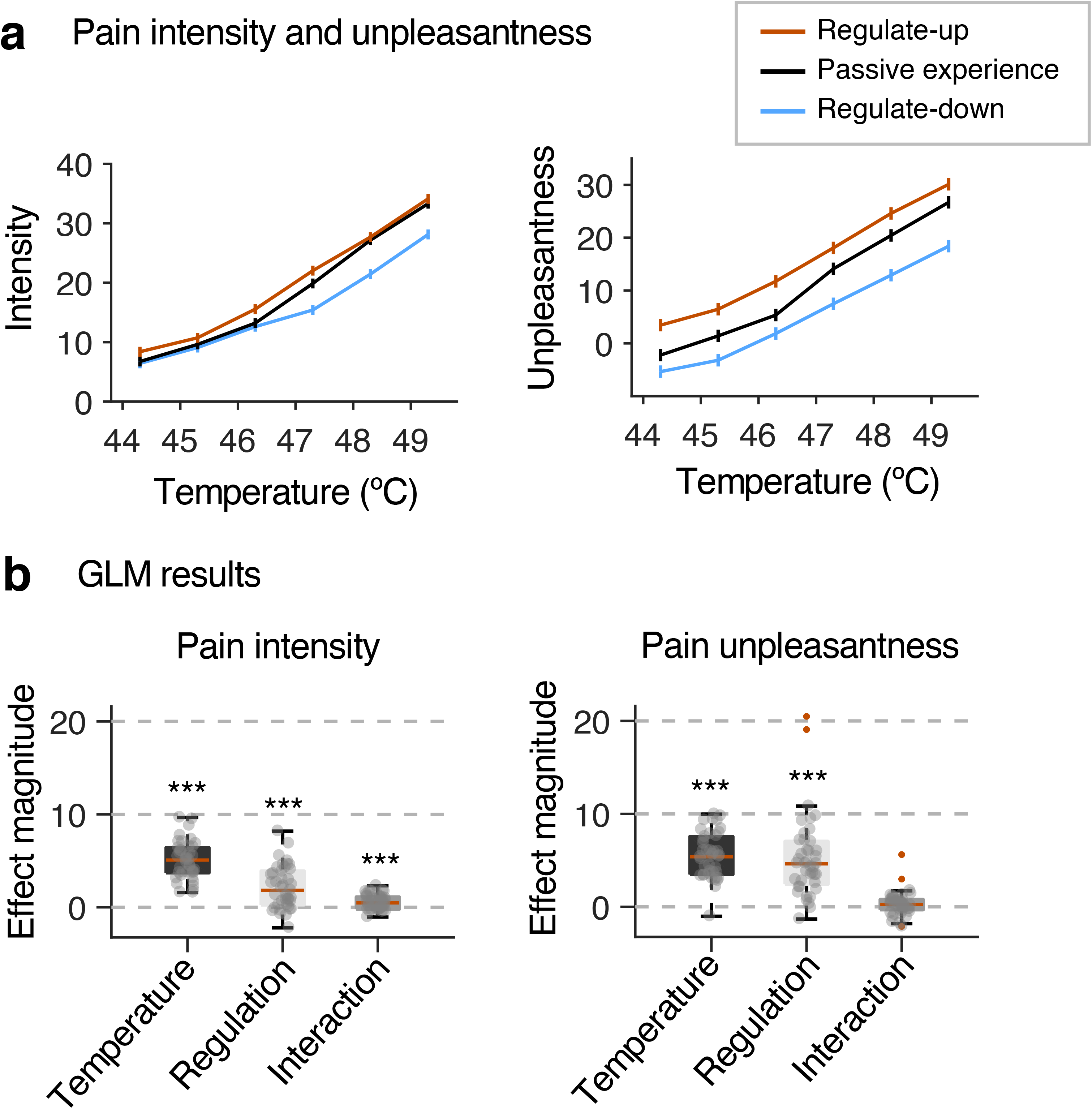
Effects of cognitive self-regulation on pain ratings. **(a)** Mean intensity and unpleasantness ratings for each temperature in the Regulate-up (red), Passive experience (black), and Regulate-down conditions (blue). Error bars represent within-subject standard errors of the mean (S.E.M.). For pain ratings, we used general Labeled Magnitude Scale (gLMS) [3]. **(b)** Effect magnitude (y-axis) represents regression coefficients (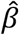) from a multi-level general linear model. Each dot shows each individual’s regression coefficient. The GLM analyses revealed that temperature (stimulus intensity, °C) and regulation (coded regulate-up, passive experience, and regulate-down as 1, 0, and −1) had significant main effects on both pain intensity and unpleasantness ratings. In addition, a significant interaction was found between temperature and regulation for the pain intensity ratings, but not for unpleasantness ratings. \*\*\**p* < .001; Bootstrap tests (10,000 iterations) were used for significance testing.

### Data acquisition

#### Electrocardiogram (ECG)

ECG activity was recorded using two 11-mm Ag/AgCl electrodes (Biopac systems, Goleta, CA) placed on the right clavicle and left lower rib area, and sampled at 500 Hz. A maximal overlap discrete wavelet transform (modwt.m, available in the MATLAB wavelet toolbox) was used to enhance ECG signal relevant to the QRS complex, and local maximums corresponding to the R-peak of the ECG signal were isolated using the findpeaks function (findpeaks.m) of the MATLAB signal processing toolbox. Peaks were then checked manually to identify and remove outliers. Inter-beat intervals (IBI) were then calculated based on differences between adjacent peaks (See Fig. S3a).

#### Skin conductance response (SCR)

SCR activity was recorded using 11-mm Ag/AgCl electrodes (Biopac systems, Goleta, CA) attached to the medial phalanges of the middle and ring fingers of the left hand. Data was sampled at 500 Hz in Study 1, and at 1000 Hz in Study 2. The difference in sampling rate between Study 1 and 2 are not expected to affect our findings because the signal of interest and other noise components are located at much lower frequencies.

### Data analysis

#### Preprocessing

Physiological data (SCR activity and ECG-IBI time-series data) was put through a low-pass filter, 5 Hz for SCR [24], and 1 Hz for ECG-IBI [14], to remove noise, and then downsampled to 25 Hz (See Fig. S3a).

#### Grand average

For each trial, a baseline was created by averaging physiological time-series data from 3 seconds before the thermal stimulation onset. A stimulus-locked physiological response was generated by subtracting the baseline value from the data in the 20-second period after the stimulation onset (See Fig. S3b). The stimulus-locked physiological responses were averaged across regulation conditions to create a mean physiological response for each temperature (Figs. 2a-b).

#### Physiological pain marker development (Analysis 1)

To develop SCR and ECG markers for pain, we first created features by averaging the stimulus-locked physiological responses in only the *passive experience* runs. This resulted in a 6 (temperature levels) × 500 (25 Hz × 20 seconds) average time-series matrix for each participant. Mean pain ratings for each participant corresponding to the six temperatures were made into a 6 × 1 vector. These data were then concatenated across participants and used for subsequent modeling (see Fig. S3 for more details). Then, principal component regression (PCR) was used to create a SCR and ECG time-course model predictive of pain ratings [20]. We chose to use PCR because it works well with the data in which the number of features (or predictors) is greater than the number of observations and the features are intercorrelated. The PCR was achieved in two steps: First, principal component analysis was conducted to reduce dimensions of features using covariance information among SCR and ECG time-series data; Second, multiple linear regression was conducted on the component space (i.e., using component scores) to predict pain ratings. In this step, we used a reduced number of components (2-3 components depending on the models) based on a leave-one-participant-out cross-validation procedure (see Fig. S4 for details). The regression model was then projected to the time space, yielding a time-series pattern of predictive weights. Bootstrap tests were conducted to identify which time points reliably contributed to the prediction [12]. For each iteration of bootstrap tests, we randomly resampled participants with replacement and trained PCR on each resampled dataset. After 10,000 iterations, we calculated the *p*-values based on the sampling distribution of predictive weights. For the correction for multiple comparisons, we used false discovery rate (FDR) *q* < .05.

**Figure 3.**
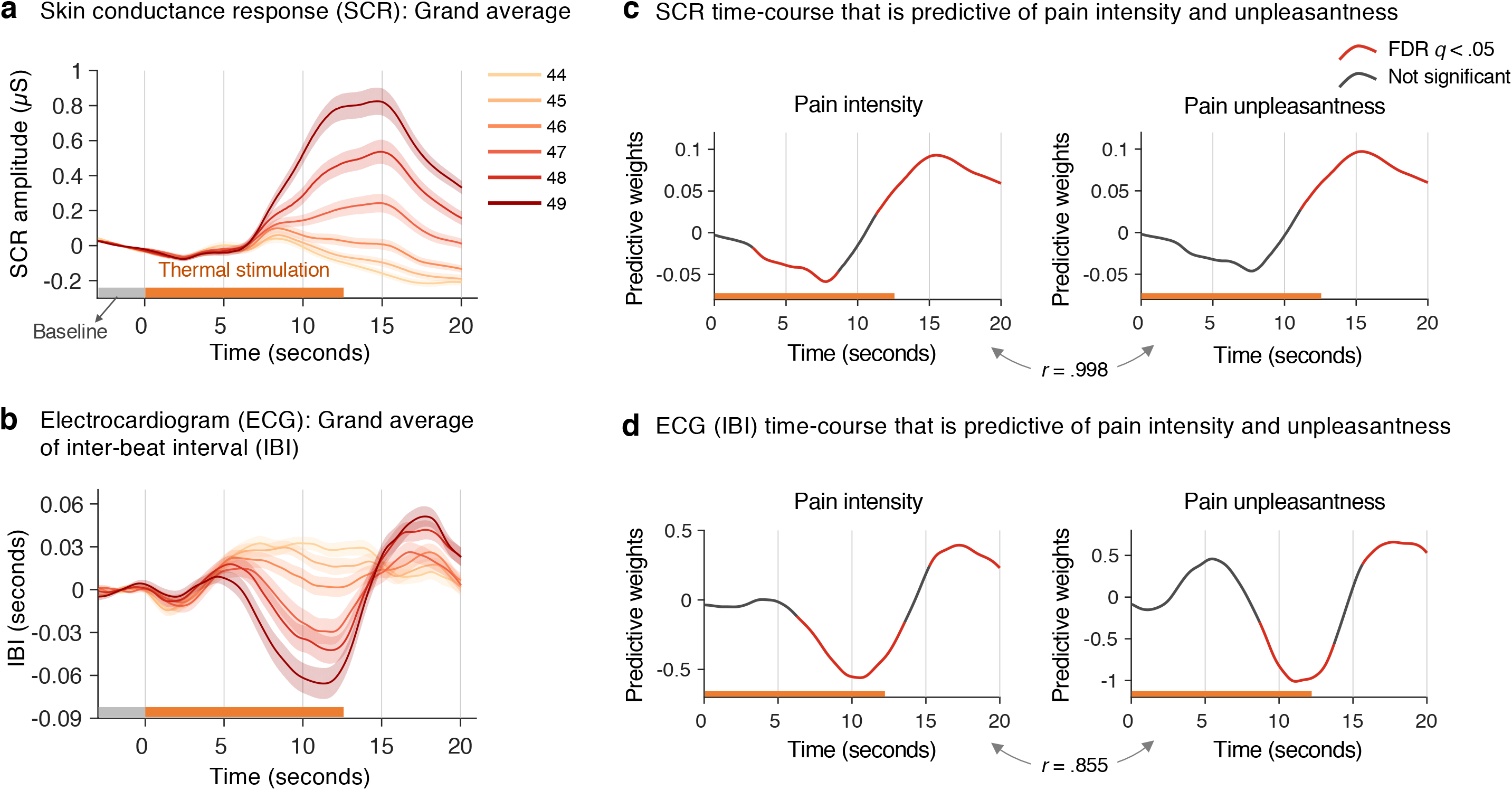
SCR and ECG’s IBI time-courses predictive of pain ratings. **(a)** Stimulus-locked grand average of skin conductance responses (SCR) across participants for each temperature. Data from 3 seconds prior to the thermal stimulation onset were used as a baseline (see **Methods** for details). Shading represents S.E.M. **(b)** Grand average of inter-beat interval (IBI) calculated from electrocardiogram (ECG). **(c)** SCR time-course markers most predictive of pain ratings (left: pain intensity, right: pain unpleasantness). We identified these markers using principal component regression based on data from passive experience runs. Regions in red represent time points that provided significantly reliable contributions to the prediction from bootstrap tests (10,000 iterations) at *q* < .05, false discovery rate (FDR). SCR time courses for pain intensity and unpleasantness were almost identical, *r* = .998. **(d)** ECG (IBI) time-course markers most predictive of pain ratings (left: pain intensity, right: pain unpleasantness). The correlation between ECG time courses for pain intensity and unpleasantness was also very high, *r* = .855.

#### Testing the physiological marker (Analysis 2 and 3)

For testing the marker on **Study 1**’s regulation data, we used a leave-one-participant-out cross-validation procedure. The time-series weights predictive of pain were derived based on physiological data from passive experience conditions for all participants except for one out-of-sample participant. These weights were then tested on the out-of-sample participant’s data in all three conditions by calculating the dot-product between the time-series weights and stimulus-locked physiological data. This process was done iteratively for each participant. Note that the data from regulation runs were not included in the model developing procedure at all. For testing the marker on **Study 2** data, which is completely independent from the model developing procedure, we calculated the dot-product between the time-series weights and stimulus-locked physiological data.

## Results

### Behavioral results of cognitive self-regulation

As shown in Fig. 2, we found that both the stimulus intensity of noxious heat and cognitive self-regulation strongly modulated ratings of both pain intensity and unpleasantness, replicating and extending previous findings [52]. Stimulus intensity had similar effects on ratings of both pain intensity and unpleasantness (intensity ratings: 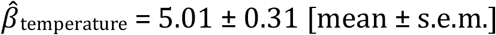, *z* = 3.86, *p* < .001 in a bootstrap test with 10,000 times resampling; unpleasantness ratings: 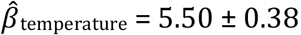, *z* = 3.71, *p* < .001). Self-regulation to increase vs. decrease pain influenced both intensity and unpleasantness ratings in accordance with regulatory goals, but influenced pain unpleasantness more strongly than intensity (unpleasantness: 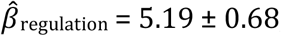, *z* = 4.54, *p* < .0001; intensity: 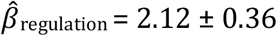, *z* = 3.94, *p* < .0001). The self-regulation effects on pain unpleasantness ratings were comparable in magnitude to a 1°C change in heat stimulus intensity, 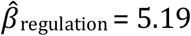 vs. 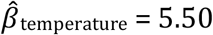. Self-regulation effects on pain intensity were larger for more intense stimuli, as evidenced by a small but significant stimulus intensity × regulation condition interaction, 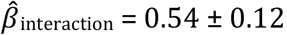, *z* = 3.81, *p* < .001. However, we found only marginal interaction effects for pain unpleasantness ratings, 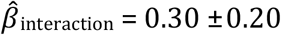, *z* = 1.82, *p* = .069. Significant modulation effects were also observed when regulate-up and regulate-down were separately compared with passive experience (all *p* values < .01 for both intensity and unpleasantness ratings; please see Table S1 for results and statistics).

### Autonomic effects of cognitive self-regulation without isolating the pain-related component

When the stimulus-locked SCR and ECG data (20 seconds after stimulus onset) were averaged for each temperature level, we observed reliable stimulus intensity-related increases in SCR amplitude and heart rate (Figs. 3a-b). In addition, when the SCR and ECG data were averaged within each regulation condition and compared, we observed small increases and decreases in SCR amplitude and heart rate for regulate-up and regulate-down, respectively (Fig. S5). Regulation-induced physiological changes were marginally significant or non-significant; for example, when using baseline-to-peak amplitudes for regulate-up vs. regulate-down, SCR: 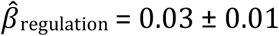, *z* = 1.93, *p* = .053, ECG: 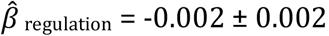, *z* = −0.63, *p* = .529; when using the area-under-the-curve, SCR: 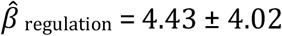, *z* = 1.14, *p* = .254, ECG: 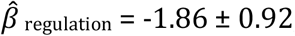, *z* = −2.03, *p* = .043. These summary measures do not, however, permit a test of whether cognitive self-regulation impacts *pain-related* physiology, due to potential masking by physiological responses to cognitive regulation demand itself, as discussed above.

### Analysis 1: Developing pain-predictive physiology markers based on SCR and ECG temporal dynamics (Study 1)

To examine autonomic changes more directly linked to pain, we first developed pain-predictive SCR and ECG markers using data from passive experience runs (i.e., pain without regulation). We then tested these markers on data from the regulation runs using a leave-one-participant-out cross-validation procedure.

As shown in Figs. 3c-d, the bootstrap test results showed that, for the SCR model, the time points between 2.7 and 8.6 seconds and between 11.3 and 20 seconds after the heat onsets were reliable predictors of pain intensity across participants (*q* < 0.05 False Discovery Rate corrected), and the time points between 11.1 and 20 seconds made reliable contributions to the prediction of pain unpleasantness. For the ECG model, the time points between 6.3 and 13.5 seconds and between 15.2 and 20 seconds were reliable predictors of pain intensity ratings, and the time points between 8.7 and 13.7 seconds and between 15.6 and 20 seconds were reliable predictors of pain unpleasantness ratings.

Cross-validated test results on the held-out participants’ data from passive experience runs showed that the mean within-participant correlations (across averaged trial responses for each stimulus intensity) of predicted with observed pain were *r* = .83 ± 0.025, *p* < .0001 (based on a bootstrap test with 10,000 resamples) for the SCR pain intensity model and *r* = .76 ± 0.047, *p* < .0001 for the SCR unpleasantness model. For ECG models, the mean prediction-outcome correlations were *r* = .60 ± 0.073, *p* < .0001 for the pain intensity model and *r* = .55 ± 0.083, *p* < .0001 for the pain unpleasantness model (Fig. S4). Thus, both SCR and ECG reliably predicted within-person variation in pain reports across trials.

We then tested whether these markers predicted pain reports during regulation runs, using cross-validation to apply the models only to new (held-out) participants (Fig. 4). Because the data from held-out participants’ regulation runs were never included in the model development process, they provided an unbiased test of whether the SCR and ECG models are predictive of pain ratings in this sample. The mean prediction-outcome correlations were *r* = .82 ± 0.020, *p* < .0001 and *r* = .73 ± 0.039, *p* < .0001 for SCR pain intensity and unpleasantness models, and *r* = .67 ± 0.045, *p* < .0001 and *r* = .65 ± 0.046, *p* < .0001 for ECG pain intensity and unpleasantness models, respectively. Thus, the correlations between autonomic responses and pain reports are similar for all regulation conditions. We address effects of regulation on the amplitude of marker responses in Analysis 3, below.

**Figure 4.**
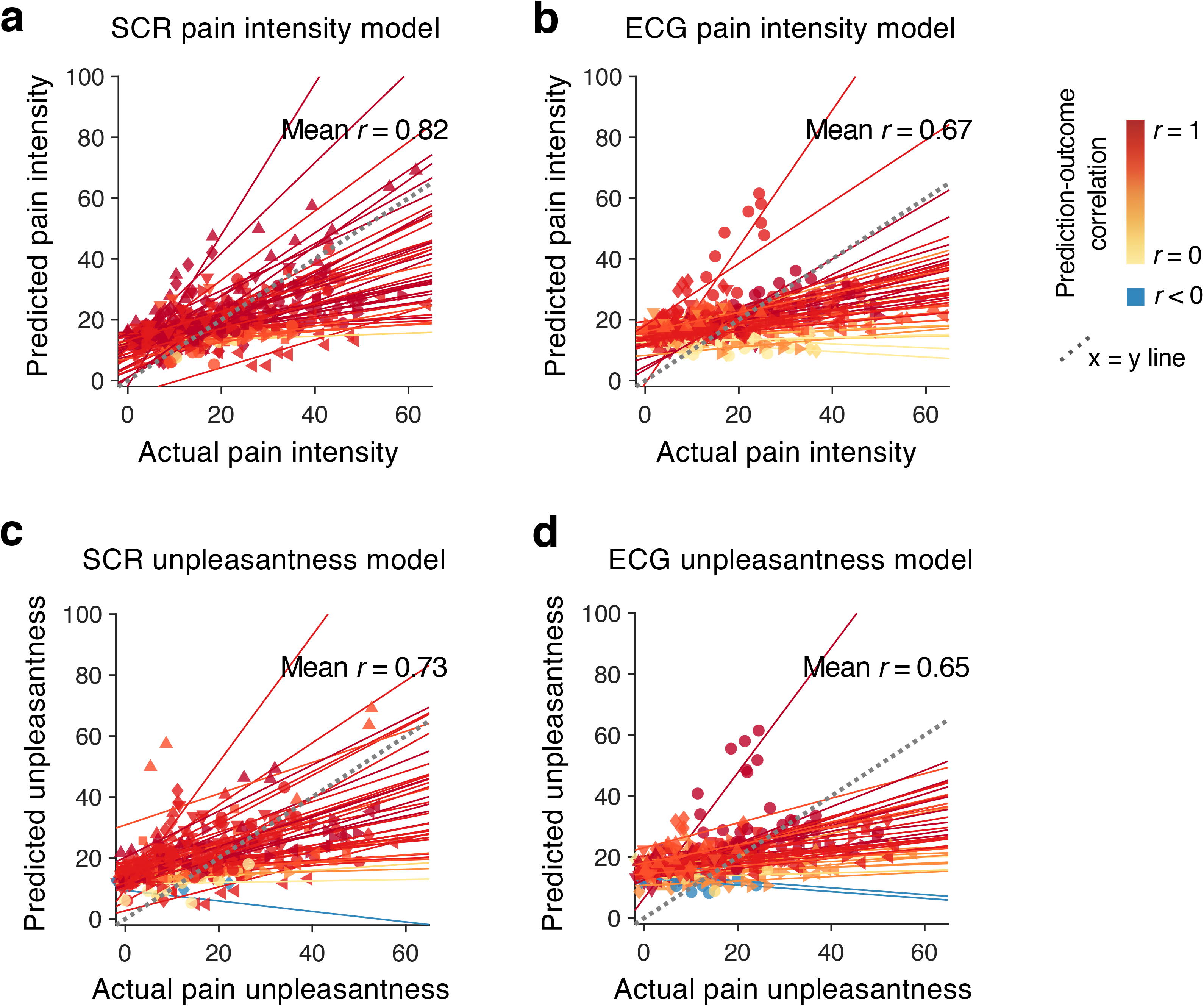
Testing the predictive models on held-out participants’ regulation data. Test results of the SCR and ECG markers on data from regulation runs (12 trials per person). Note that no data from regulation runs were used for marker development. Additionally, leave-one-participant-out cross validation was used to prevent any possibility of overestimation of model performance due to dependency among data from same individuals. Actual pain intensity (**a** and **b**) or actual pain unpleasantness (**c** and **d**) versus cross-validated predicted pain intensity or unpleasantness are shown in the plots, and each line or symbol represents individual participant’s data. The line color represents a correlation level for each participant (red: higher *r*; yellow: lower *r,* blue: *r* < 0), and the dotted line represents points where the actual pain ratings are same with the predicted ratings (i.e., x = y).

### Analysis 2: Testing the SCR marker on an independent dataset (Study 2)

Grand averages and baseline-to-peak amplitudes of stimulus-locked SCR showed enhanced responses to increasing stimulus intensity in both pain receivers and their partners who observed pain (Figs. 5a-c). As shown in Fig. 5c, the baseline-to-peak SCR amplitude significantly increased for 49 vs. 47°C and 48 vs. 47°C in both pain receivers (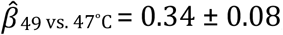, *z* = 4.98, *p* < .0001, 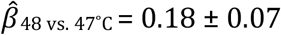, *z* = 5.76, *p* < .0001) and pain observers (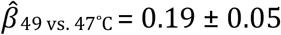, *z* = 4.79, *p* < .0001, 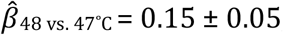, *z* = 4.12, *p* < .0001). For 49 vs. 48°C, the baseline-to-peak amplitude showed significant increases only in pain receivers, 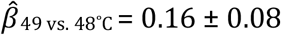, *z* = 2.83, *p* = .0046, not pain observers, 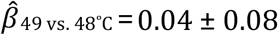, *z* = 0.51, *p* = .608. In addition, experiencing pain induced larger overall SCR changes than observing pain, 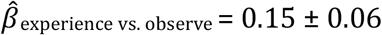, *z* = 2.66, *p* = .0077. Thus, standard SCR amplitudes showed significant increases proportional to stimulus intensity for both experienced and observed pain, and limited selectivity for pain experience.

**Figure 5.**
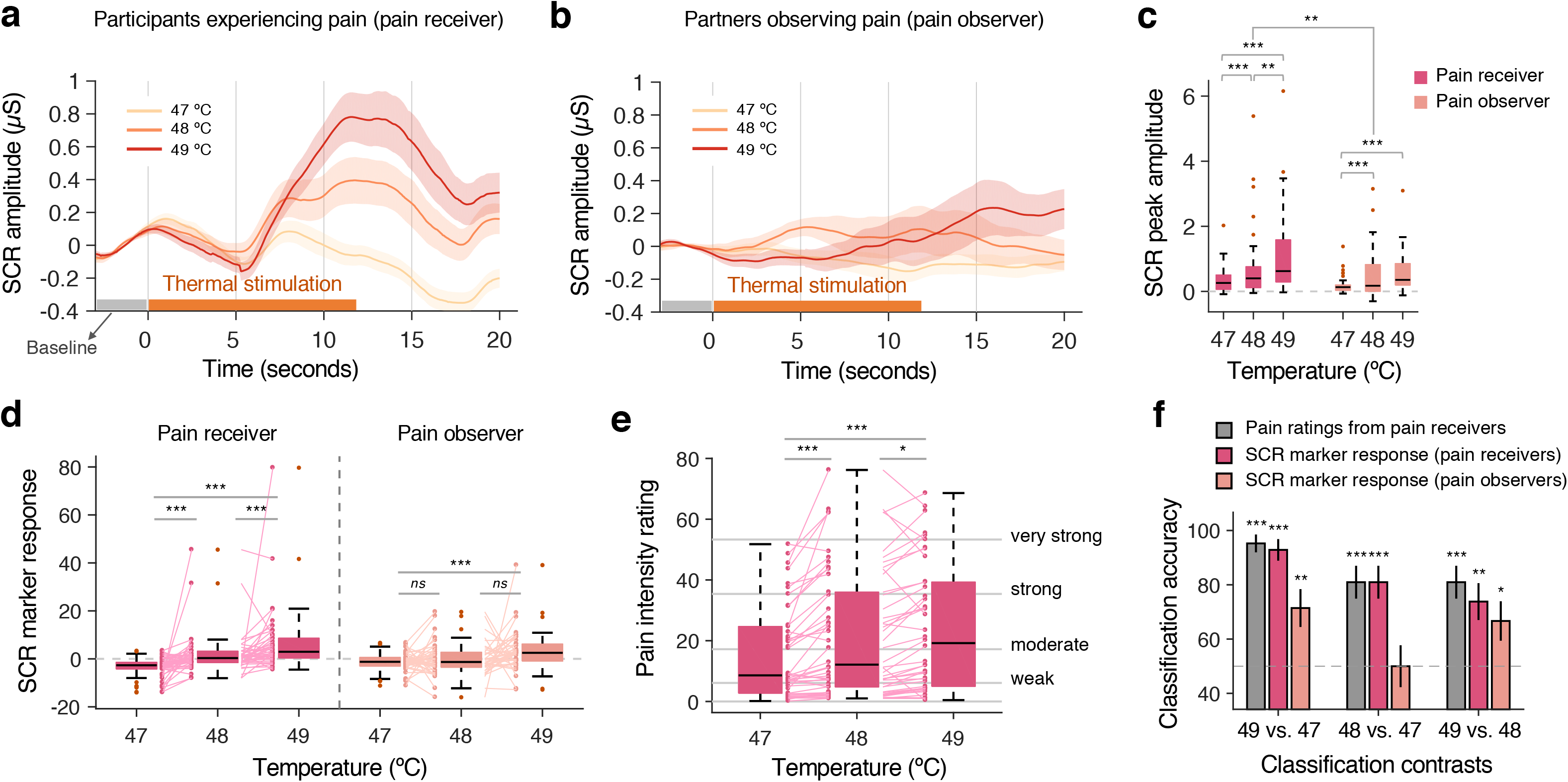
Validation of markers on an independent data set. We tested the sensitivity and specificity of the SCR pain intensity marker using data from Study 2. In Study 2, participants received thermal heat stimulations on their leg (pain receiver), and their romantic partners observed the main participants experiencing pain (pain observer). **(a)** Mean SCR amplitudes of pain receivers during three different stimulation temperatures. Shading represents S.E.M. **(b)** Mean SCR amplitudes of pain observers during the same trials. **(c)** Baseline-to-peak SCR amplitudes from pain receivers and observers for three temperature levels. **(d)** SCR marker responses from pain receivers and observers for three temperature levels. Lines connect the same individuals’ marker responses. **(e)** Mean pain intensity ratings from pain receivers for three temperatures. Lines connect the same individuals’ pain ratings. **(f)** The two-choice classification accuracy for stimulus intensity contrasts using (i) pain ratings from pain receivers, (ii) SCR marker responses of pain receivers, and (iii) SCR marker responses of pain observers. \*\*\**p* < .001, \*\**p* < .001, \**p* < .05 (two-tailed); Bootstrap (c, d, and e) and binomial test (f) was used for significance testing.

When we tested the SCR pain intensity marker from Analysis 1, the SCR marker tracked the changes in first-person pain better than in observed pain, demonstrating the marker’s differential sensitivity and specificity to first-person pain. The SCR marker showed significant increases for 48 vs. 47°C, 49 vs. 48°C, and 49 vs. 47°C in the participants who experienced pain, 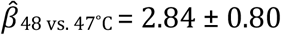, *z* = 5.58, *p* < .0001, 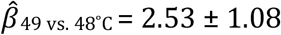, *z* = 4.71, *p* < .0001, and 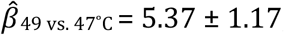, *z* = 5.44, *p* < .0001, respectively (Fig. 5d). Effect sizes for 1°C increase ranged from Cohen’s *d* = 0.73 to *d* = 0.86. These increases were comparable in magnitude to the increases in pain ratings, 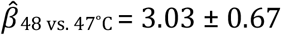, *z* = 4.60, *p* < .0001, 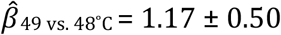, *z* = 2.07, *p* = .0386, and 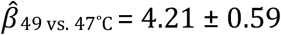, *z* = 3.74, *p* < .001 (Fig. 5e). Effect sizes for 1°C increase on pain ratings were *d* = 0.32 to *d* = 0.71. Conversely, during observed pain, the SCR marker showed non-significant or marginal increases for 48 vs. 47°C and 49 vs. 48°C, 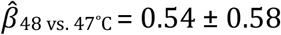, *z* = 0.98, *p* = .33, 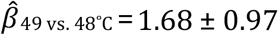, *z* = 1.94, *p* = .052 (Fig. 5d). The SCR marker did show a significant, but relatively small, increase for 49 vs. 47°C during observed pain, 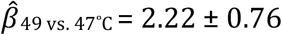, *z* = 3.58, *p* < .001. The effect sizes for observed 1°C increase were *d* = 0.15 and *d* = 0.30, respectively, approximately 3 times lower than those for first-person pain. Overall, our SCR marker showed strong increases for pain experience and weak responses for pain observation, demonstrating good sensitivity, but a moderate level of specificity in this context.

To further characterize stimulus intensity-related increases in the SCR marker, we conducted pairwise classification tests, in which pain ratings and SCR marker responses between two levels of stimulus intensity (i.e., 49 vs. 47°C, 48 vs. 47°C, and 49 vs. 48°C) were compared, and the condition with the higher levels of pain rating or SCR marker response was predicted to be the more intense (Fig. 5f). During somatic pain experience, the SCR marker demonstrated high accuracy in forced-choice discrimination of different levels of stimulus intensity; for 49 vs. 47 °C, accuracy = 92.9% ± 4.0, *p* < .0001, for 48 vs. 47 °C, accuracy = 81.0% ± 6.1, *p* < .0001, for 48 vs. 49 °C, accuracy = 73.8% ± 6.8, *p* = .0029. These results were comparable to the performance obtained when using self-reported pain to predict which condition had a more intense stimulus: for 49 vs. 47 °C, accuracy = 95.2% ± 3.3, *p* < .0001, for 48 vs. 47 °C accuracy = 81.0% ± 6.1, *p* < .0001, for 49 vs. 48 °C = 81.0% ± 6.1, *p* < .0001. For observed pain, the marker response showed worse classification performance than the response to somatic pain, for 49 vs. 47 °C, accuracy = 71.4% ± 7.0, *p* = .0079, for 48 vs. 47 °C accuracy = 50.0% ± 7.7, *p* = 1.00, for 49 vs. 48 °C = 66.7% ± 7.3, *p* = .0436. Though some of the contrasts were significantly above the chance level, if we corrected these test results for multiple comparisons (9 tests in this classification) using a Bonferroni method (i.e., corrected $ = 0.05/9 = 0.0056), all the classification results for observed pain became non-significant, while all first-person pain results remained significant.

### Analysis 3: The effects of cognitive self-regulation on the pain-predictive physiology markers (Study 1)

To test whether cognitive self-regulation has significant impacts on pain-related physiology, we conducted multi-level general linear models using the SCR and ECG marker response calculated from Study 1 data as outcome measures and tested the effects of stimulus intensity, self-regulation (regulate up vs. down), and their interaction.

Both stimulus intensity and self-regulation had significant effects on the SCR and ECG pain intensity and unpleasantness markers (Fig. 6 and Fig. S6); for the SCR intensity marker, 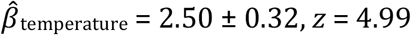, *z* = 4.99, *p* < .0001, 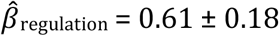, *z* = 4.34, *p* < .0001, for the SCR unpleasantness marker, 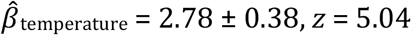, *z* = 5.04, *p* < .0001, 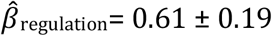, *z* = 4.11, *p* < .0001, for the ECG intensity marker, 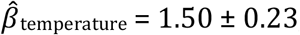, *z* = 4.95, *p* < .0001, 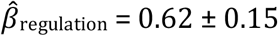, *z* = 4.16, *p* < .0001, and for the ECG unpleasantness marker, 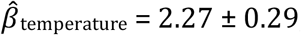, *z* = 4.13, *p* < .0001, 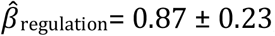, *z* = 4.09, *p* < .0001. Regulation effect sizes (up vs. down) were in the “moderate to large” range, between *d* = 0.63 and *d* = 0.67 for all models.

**Figure 6.**
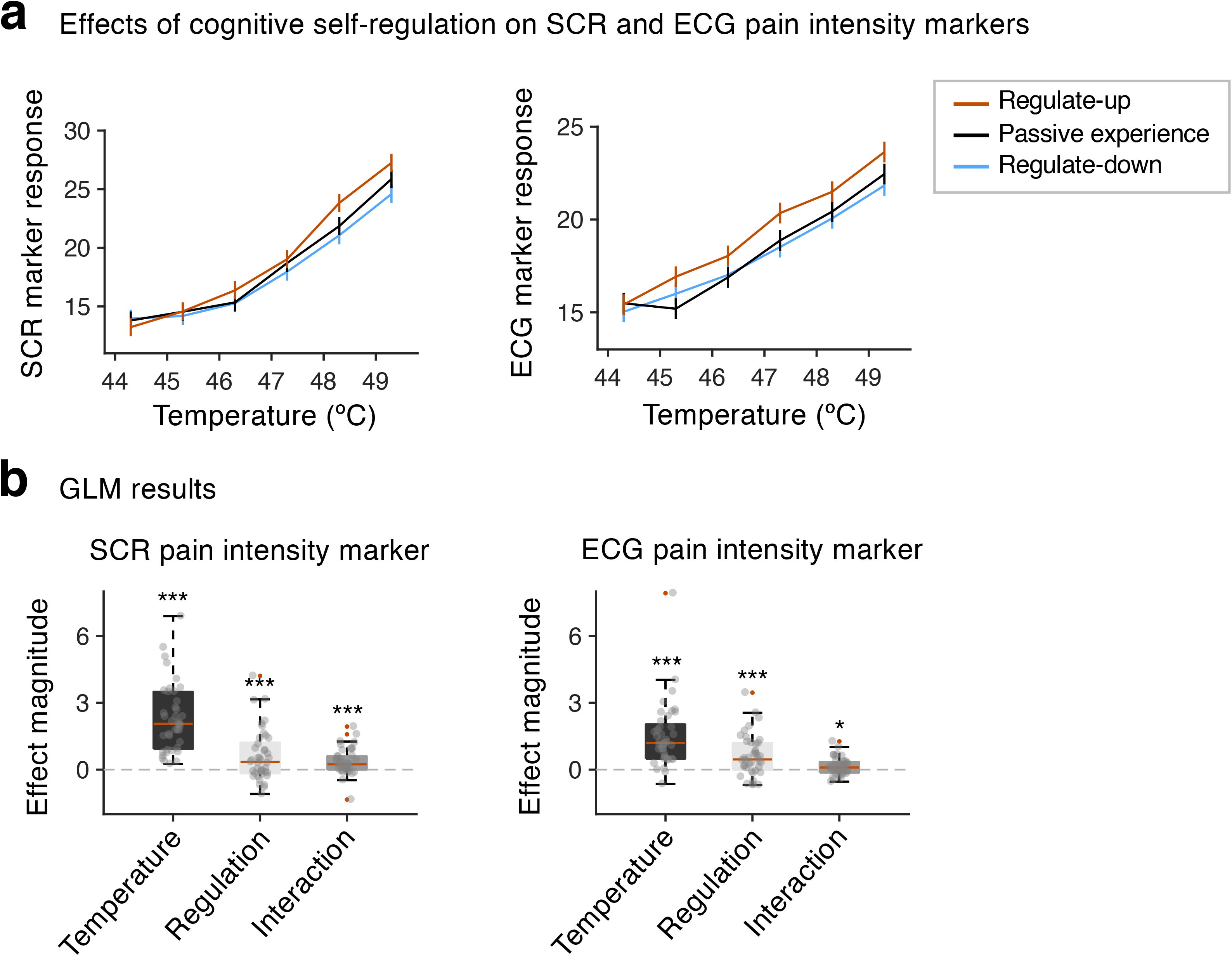
Effects of cognitive self-regulation on SCR and ECG markers. **(a)** This is an analogous plot to Fig. 1a’s left panel, but here we used predicted pain scores based on SCR and ECG pain intensity models. Error bars represent within-subject S.E.M. **(b)** Multi-level general linear model results. Similar to the behavioral findings, both stimulus intensity and cognitive self-regulation had significant effects on SCR and ECG marker responses, indicating that cognitive self-regulation has significant effects on pain-related autonomic physiology. \*\*\**p* < .001; Bootstrap tests (10,000 iterations) were used for significance testing.

Similar to the results with pain intensity ratings, the effects of self-regulation on SCR and ECG pain intensity markers showed significant interactions with stimulus intensity, 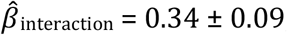, *z* = 3.90, *p* < .0001, and for the ECG intensity marker, 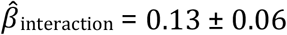, *z* = 2.43, *p* = .015, suggesting that the self-regulation effects on pain intensity-related physiology increase as stimulus intensity increases. For the pain unpleasantness markers, a significant interaction was observed in SCR, 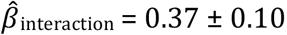 ± 0.10, *z* = 3.65, *p* < .001, but not in ECG, 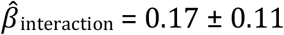, *z* = 1.71, *p* = .087.

### Estimating regulation effect sizes in terms of effective changes in stimulus intensity

We standardized the beta coefficients of self-regulation relative to those of stimulus temperature, in order to compare the effect magnitudes of self-regulation on different outcome variables. We used the effect size of a 1°C change in temperature as a reference; for example, the relative effect magnitude of 0.42 for the effects of self-regulation on pain intensity ratings and 0.94 on pain unpleasantness ratings are comparable to the effects of a 0.42 °C and 0.94 °C change in temperature on pain intensity and unpleasantness, respectively. As shown in Fig. 7, the self-regulation effects on ECG markers were larger in magnitude than the regulation effects on SCR markers and were comparable to the effects on pain intensity ratings; relative effect magnitude for SCR intensity marker = 0.24 °C, SCR unpleasantness marker = 0.22°C, ECG intensity marker = 0.41°C, and ECG unpleasantness marker = 0.38°C. Thus, self-regulation has the largest effects on pain unpleasantness (0.94 °C), followed by pain intensity (0.42 °C) and heart rate (0.38-0.41 °C), followed by SCR (0.22-0.24 °C).

**Figure 7.**
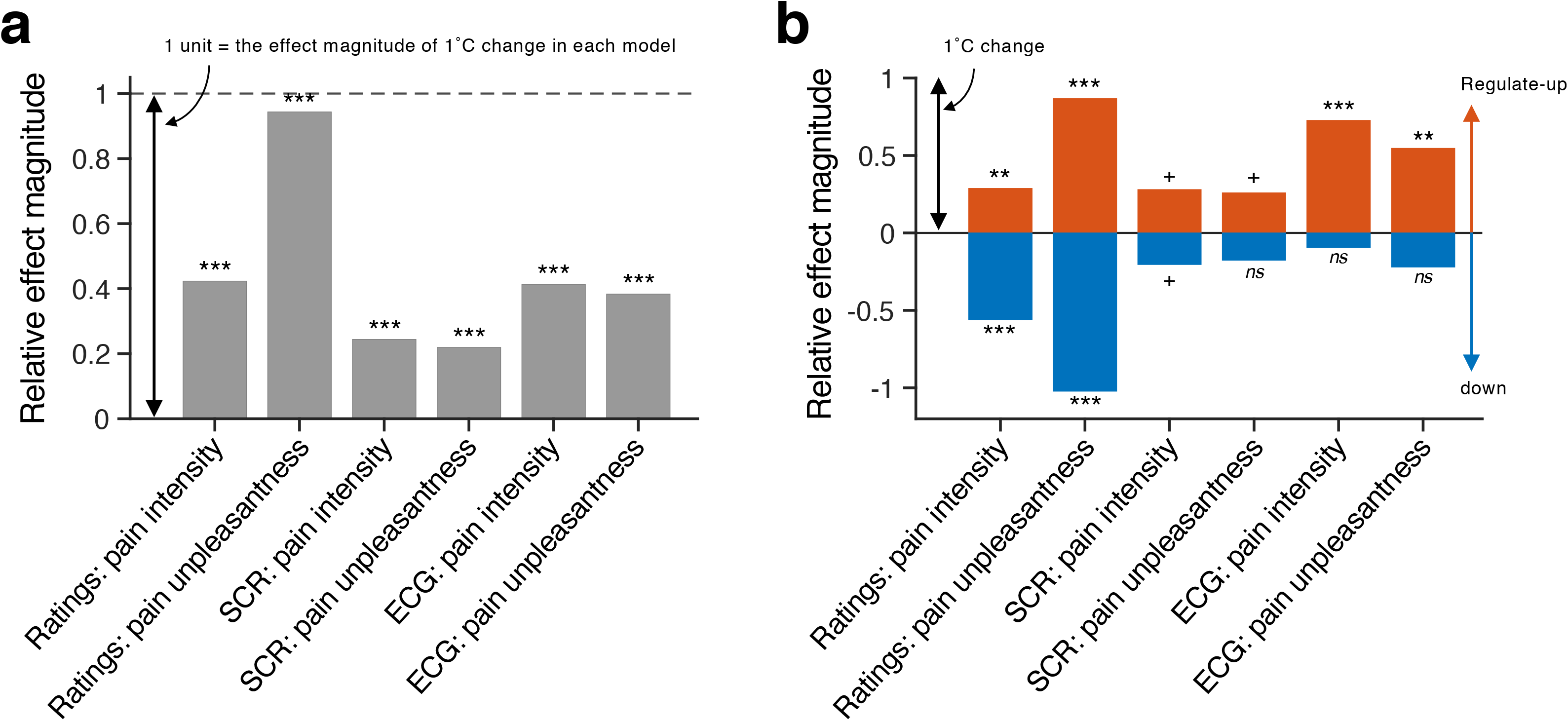
Relative effect magnitudes for different outcome variables. To compare the effect magnitudes across different models, we calculated relative effect magnitudes of self-regulation on different outcome measures by comparing them to the effects of stimulus intensity. In other words, in both of these plots, 1 unit in the y-axis indicates the effect magnitude comparable to the effects of a 1°C change in stimulus intensity. The x-axis shows the various outcome measures assessed in the study; SCR- and ECG-related measures are responses in trained, pain-predictive models. **(a)** Relative effect magnitude for the average changes by regulate-up vs. regulate down. For example, the relative effect magnitude of 0.94 for pain unpleasantness ratings can be interpreted that regulate-up and regulate-down on average had effects on pain unpleasantness comparable to the effects of 0.94 °C change in stimulus intensity. **(b)** Relative effect magnitude separately for each regulation direction. The significance test results were from bootstrap tests (10,000 iterations) of general linear models for different outcome variables. *^ns^ p* > .1, ^+^*p* < .1, \*\**p* < .01, \*\*\**p* < .001.

Unlike the effects on pain ratings, the self-regulation effects on physiological markers seem largely driven by the regulate-up condition rather than the regulate-down condition, but the differences in beta coefficients between regulate-up vs. passive experience and passive experience vs. regulate-down were not significant; for the SCR intensity marker, mean difference 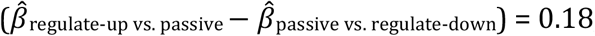, *t*_40_ = 0.30, *p* = 0.762, for the SCR unpleasantness marker, mean difference = 0.23, *t*_40_ = 0.34, *p* = 0.732, for the ECG intensity marker, mean difference = 0.95, *t*_39_ = 1.74, *p* = 0.090, for the ECG unpleasantness marker, mean difference = 0.74, *t*_40_ = 0.95, *p* = 0.340 (Table S1).

In sum, increasing the intensity of painful stimuli had a strong effect on autonomic responses, allowing us to identify a pain-predictive temporal waveform in both ECG and SCR that can be applied to new individuals and studies to evaluate a pain-related component of responses to noxious stimuli. Cognitive self-regulation significantly modulated these pain-predictive autonomic markers, with effects that appeared strongest for up-regulation of pain.

## Discussion

Although the effects of cognitive interventions on self-reported pain are well-documented [15], their effects on autonomic physiology are less clear. A historical problem with assessing autonomic effects is that they reflect a mix of components related to orienting, arousal, and pain. Here, we identified a pain-related component of autonomic responses to noxious stimuli that is distinct from the overall SCR and heart-rate responses to noxious events. The component waveform involves early decreases and late increases just post-stimulus-offset, effectively subtracting late from early activity during stimulation. This late activity occurs just after peak reported pain, which peaks at the end of the stimulation period in previous studies [28; 48]. Thus, it is less sensitive to the autonomic responses driven by novel stimulus onset and orienting [17; 50], and more selective for pain. This waveform can be thought of as a link function that averages autonomic activity over a painful stimulation period into a single value optimized to track post-trial pain reports. These functions—one for each of ECG (heart-rate) and SCR responses linked to each of pain intensity and affect—can be applied to new individuals to generate testable predictions about pain based on autonomic responses. Turning back to the question of self-regulation, we did not find cognitive regulation effects on standard baseline-to-peak amplitude or area-under-the-curve measures of event-related autonomic responses. However, when we applied the waveforms to isolate a pain-related component of autonomic activity, we observed significant cognitive regulation effects on both heart rate and SCR with meaningful effect sizes. These findings suggest that cognitive strategies have effects on pain-related aspects of autonomic function.

An important aspect of this study is that we developed SCR and ECG physiological markers for pain first, and then applied these markers to examine the effects of cognitive pain modulation on pain-related physiology. These markers can be used in future studies to test relationships with pain and influences of multiple types of interventions. The physiological markers developed here have reasonable levels of sensitivity, specificity, and generalizability in predicting pain across two independent datasets. Tests in out-of-sample individuals showed strong correlations with pain reports (*r* = 0.55-0.83) and showed the ability to track pain and differentiate first-person pain experience from observation of another person in pain, an experience that activates the autonomic nervous system, but with a different temporal profile. However, testing sensitivity and specificity should be an open-ended process [51]. The current study provided only a limited set of tests for sensitivity and specificity of the markers, and thus further validation with different experimental conditions will be required to precisely characterize them. Despite these challenges, these predictive models have the potential to provide an additional, cost-effective way to objectively assess acute pain besides existing neuroimaging-based pain markers (e.g., ref. [48]). In addition, these markers allowed us to examine the effects of self-regulation on pain-related physiology in a more specific manner by isolating pain-related autonomic response from some other non-specific factors that are present in physiological measurements. Importantly, when we tested an SCR marker from a previous study by Geuter et al. [18] on our study dataset, the SCR marker showed similar predictive performances predicting pain ratings and similar responses to cognitive self-regulation, despite the differences in stimulus duration (Fig. S7). This suggests that the physiological markers could be robust to some changes in stimulus parameters, though future studies could develop more generalizable models across types of stimulation.

Our analysis results revealed some interesting patterns in physiological responses to pain and pain modulation, though we will need additional studies to confirm these to be robust and reproducible. First, the SCR and ECG time points that reliably contributed to the prediction of pain intensity occurred earlier than the time points predictive of pain unpleasantness (Figs. 3c-d). This suggests that the sensory and discriminative processes that are closely related to pain intensity may precede the generation of pain unpleasantness [37; 38]. Second, when comparing the magnitude of self-regulation effects to those of varying stimulus intensity, self-regulation has stronger effects on unpleasantness than intensity, and stronger effects on cardiovascular responses (ECG) than responses in the skin (SCR) (Fig. 7a). For example, the effect of self-regulation on the ECG pain intensity marker was comparable to a 0.41 °C change in stimulus intensity, whereas the SCR pain intensity marker showed a regulation effect comparable to a 0.23 °C change.

The markers for pain unpleasantness showed a similar pattern. An interesting observation from our findings is that the regulate-down condition showed stronger effects on pain ratings as temperature increased, while the regulate-up condition showed weaker effects on pain ratings as temperature increased. While we did not directly assess motivation or beliefs about regulation, this finding suggests that motivation to modulate pain may be an important factor in its efficacy.

Another interesting observation was that the effects of self-regulation on pain-related physiology seem to be largely driven by the regulate-up condition rather than the regulate-down condition, especially for the ECG markers (Fig. 7b). When tested individually against the passive experience condition, the regulate-up condition showed larger effect magnitudes for the SCR and ECG physiological marker responses, a trend not seen for pain ratings. This finding may support for the asymmetric effects of up vs. down-regulation on pain-related physiology, but direct comparisons between beta coefficients for regulate-up and -down against the passive experience condition yielded null results (all *p*s > .05). It is also possible that we have null effects for passive experience vs. regulate-down simply because of the lack of sufficient statistical power to test each regulation direction separately. We need future studies with larger numbers of trials in each condition to get definitive answers for whether asymmetrical effects of regulate-up vs. down on pain-related physiology exist or not.

This study has some additional limitations that should be addressed in future studies. First, our sample was racially homogenous (87% Caucasian), and therefore our findings must be interpreted and generalized with caution. Second, despite our effort to standardize the regulation instructions and strategies across participants (e.g., appearance of experimenter, intonation of verbal instructions, rapport before and during the experiment), we found that participants used a diverse set of regulation strategies from post-experiment questionnaires (Fig. S2). Future studies examining the effects of using different regulation strategies on pain, physiological, and neural outcomes would be very informative. Third, the SCR marker was tested for the specificity between experienced pain versus observed pain, but not with non-noxious somatosensory conditions, such as using non-noxious thermal, auditory, visual, or taste stimuli. Lastly, although we instructed participants not to regulate pain during the passive experience runs, intrinsic and spontaneous coping responses to pain or carry-over effects from previous regulation runs might influence the conditions and thus were included in our physiological markers [41]. Significant differences in pain ratings between the passive experience versus down-regulation conditions (all *p*s < .0001 for pain intensity and unpleasantness ratings; see Table S1) suggest that the influences of spontaneous pain regulation on pain during the passive experience runs were small, though they may have influenced the asymmetry between up-regulation and down-regulation effects by reducing autonomic responses under ‘neutral’ instructions to some degree. Nevertheless, examining the physiological effects of spontaneous pain regulation and the temporal dynamics of regulation effects would be important future research topics.

To conclude, in this study, we showed that cognitive self-regulation operates on the level of the autonomic nervous system, producing physiologically meaningful changes. Understanding the nature of the relationship between cognitive regulation and pain physiology has implications for the fields of both basic and clinical pain research. It can provide insight into the neurophysiological mechanisms underlying cognitive and other types of pain regulation. Additionally, our study can be useful for clinical pain management, as our regulation method shares common elements with techniques such as cognitive behavioral therapy and mindfulness- and acceptance-based therapies [25; 49]. We believe that showing that these techniques can modulate pain physiology has a powerful message for physicians and other caregivers, and for patients.

It has been a long-standing challenge for clinicians and researchers to find physiological markers for pain [29]. The predictive modeling approach used here represents a potential avenue through which quantitative biological measures related to pain can be developed and tested across studies. These methods have several potential clinical applications, but creating biomarkers for pain is especially important. Use as a surrogate biomarker in place of pain is unlikely to be viable in the near future, but the United States Food and Drug Administration (FDA) defines biomarkers for multiple other purposes (e.g., https://www.fda.gov/ucm/groups/fdagov-public/@fdagov-drugs-gen/documents/document/ucm533161.pdf). For example, biomarkers for pain are needed to show that interventions engage particular mechanistic targets and track improvements over time (‘monitoring’ and ‘pharmacodynamic/response’ biomarkers). In this case, the measures we develop can show that treatments engage brainstem generators of autonomic responses, an important part of the overall response to painful events.

## Acknowledgements

This work was supported by NIH R01DA035484, R01MH076136 (T.D.W.), and IS-R015-D1 (Institute for Basic Science), 2019R1C1C1004512 (National Research Foundation of Korea) (C.-W.W). The authors have no conflicts of interest to declare.

## Data Availability Statement

De-identified data for Study 1 and the analysis scripts for reproducing all figures and results are publicly available at https://github.com/canlab/cognitive_regulation_physiology.

## Author contributions

Study 1: CWW and TDW developed the study concept. GM, CWW, and TDW contributed to the study design. GM and CWW collected data. Study 2: MCR and TDW developed and designed the study, and MCR collected data. Study 1 and 2: GM and CWW analyzed data and drafted the manuscript, and all authors revised the manuscript and provided critical feedbacks. All authors approved the final version of the manuscript for submission.

## Supplemental Methods

### Full script for the Regulation practice (Study 1)

(Ask participant to close their eyes)

Let’s take a moment and tune in to how things are for you right now. As we’re sitting here, we can create a space where you can have control over what you’re feeling, in a way that allows you to control the pain you experience. Are you ready? To begin, focus on the sensation of your left forearm that does not hurt at all in this moment, and become aware of how it feels, of the sensations that come from that region. If you are able to bring awareness to any aspects of this experience, even for the briefest of moments, you can develop a powerful relationship with your sensations, for example, pain, and even more importantly, with your mind and body.

(Pause)

*Regulate-up*

Now, we want you to practice increasing a painful sensation by using the power of your mind. Our research indicates that one of the most effective ways to do this is by changing the meaning of the painful sensation using the power of your imagination. Take a moment and try to imagine what it might feel like if this part of your body hurt (left forearm). As if something very hot or sharp was pushing on it, perhaps like one of the stimulations you just felt. Imagine how unpleasant the pain is, for instance, how strongly you would like to remove your arm from it. You can increase pain by imagining your arm burning from something very hot being put on it, and the stinging and shooting sensations that go along with that image. As you feel the pain rise, imagine it rising faster and faster, and going higher and higher. Think of how disturbing it is to be burned, and visualize your skin sizzling, melting, and bubbling as a result of the intense heat.

(Pause)

As you’re imaging this, slowly become aware of what this feels like. Don’t worry if you had trouble imagining this, it will become much easier once you have real sensations you can manipulate. With the real painful heat, you will be able to exert your power of mind over your painful sensation.

*Regulate-down*

Take a moment to return to the sensation on your forearm. Now we want you to practice decreasing the painful sensation using the power of your mind. Imagine once again the burning, stinging, and shooting sensations that go along with strong heat being applied to your left forearm.

(Pause)

Now focus on the part of that pain that feels pleasant, like the warmth on your skin of hot clothes being taken out of the dryer. Allow the pain and heat to be carried away, flowing away from your body as if being taken downstream if you were to plunge that part of you into a cold river. Think of what it might feel like to be very cold, and have the heat on your arm warm you up. Once again, even if you were able to become aware of and control any aspects of this experience even for the briefest of moments, you have already come a long way in being able to build a powerful relationship with pain in such a way that you are able to change your experience of it. Now, open your eyes.

(open eyes)

If you found this difficult, don’t worry: this was just a practice, and it will become much easier to manipulate your experience once you have stronger sensations to work with. How was it?

### Full Script for Regulation Instructions (Study 1)

*Regulate-down.* To down-regulate pain, participants were instructed to minimize the amount of pain felt by focusing on their sensations and cognitively changing the context in which they were experienced. Full instructions are as follows:

> During this section, we are going to ask you to try to imagine as hard as you can that the thermal stimulations are less painful than they are. Focus on the part of the sensation that is pleasantly warm, like a blanket on a cold day, and the aspects of the heat that are calming, soothing, and relaxing. You can use your mind to turn down the dial of your pain sensation, much like turning down the volume dial on a stereo. As you feel the stimulation rise, let it numb your arm, so any pain you feel simply washes away. Picture your skin being very cool from walking outside on a snowy day, and focus on how comforting the stimulation feels on your arm as it warms you up. Think of how you would like to keep your arm on the heat, and visualize the powerful warmth flowing and spreading through you as it gives you energy and life.

*Regulate-up.* Instructions to up-regulate pain were designed to be similar in content to down-regulation instructions, but instead maximize the amount of pain felt. Full instructions are as follows:

> During this section, we are going to ask you to try to imagine as hard as you can that the thermal stimulations are more painful than they are. Try to focus on how unpleasant the pain is, for instance, how strongly you would like to remove your arm from it. Pay attention to the burning, stinging and shooting sensations. You can use your mind to turn up the dial of the pain, much like turning up the volume dial on a stereo. As you feel the pain rise in intensity, imagine it rising faster and faster and going higher and higher. Picture your skin being held up against a glowing hot metal or fire. Think of how disturbing it is to be burned, and visualize your skin sizzling, melting and bubbling as a result of the intense heat.

*Passive experience.* Participants were asked to focus on the fixation cross on the screen, and rate intensity and unpleasantness of pain without regulating it up or down. This condition was loosely matched in length and word count to both the regulate-up and regulate down conditions. Full instructions are as follows:

> During this section, we are going to ask you to stare at the fixation cross, and rate how intense and pleasant/unpleasant each stimulation is. Try not to regulate or change your sensation, but instead accurately rate what each sensation was like as you felt it. Focus on the fixation cross during each heat stimulation, and try and keep your eyes open and your face aligned towards the computer screen. As you feel the stimulation rise, try and sit as still as possible, and keep your eyes and face oriented towards the camera in front of you.

### Study 2 Task Design

The experiment involved five task conditions. Each condition consisted of 10 trials of painful thermal stimulation delivered in a pseudorandom order to three different sites on the main participant’s left leg. Moment-by-moment pain intensity ratings were collected from the main participant each trial. Overall pain intensity and unpleasantness ratings were collected from the main participants at the end of each condition. The first condition was a “Pre-manipulation” condition where the main participant experienced the pain stimulations alone, without their partner present. The presentation order of the next three conditions was pseudorandomized so that there were six total orders. These conditions were (a) a “Present” condition, where the partner was present but did not touch or significantly interact with the main participant, (b) a “Hand-holding” condition, where the partner held the main participant’s left hand, and (c) a “Gentle stroking” condition, where the supportive participant gently stroked the forearm of the main participant in a pleasurable and soothing way. Lastly, the main participant underwent a “Post-manipulation” condition, where they again experienced pain without their partner present. Skin conductance responses were recorded from both participants (main participants and partners) during each block they participated in. The current study uses data only from the “Present” condition.

---

**Figure S1.**
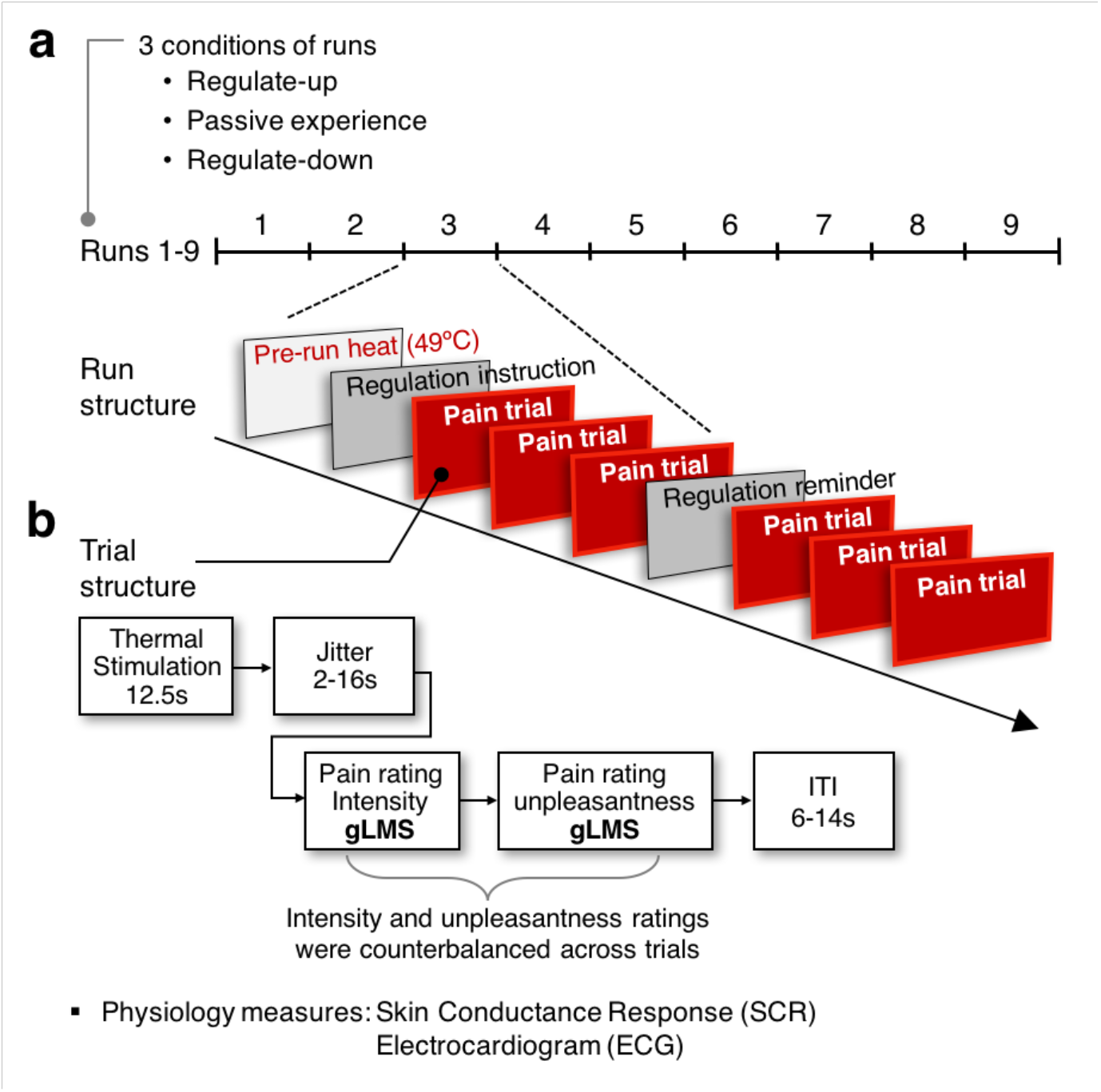
Experimental Design. **(a)** Run structure: Three different regulation conditions were pseudorandomized across 9 runs using a Latin square method. Each run began with a pre-run heat stimulation to minimize peripheral habituation effects on pain experience [3]. After instructions for regulation were presented, six trials of heat pain were administered, and a regulation reminder was given in the middle of the run. **(b)** Trial structure: Each trial consisted of a thermal stimulation of 12.5 seconds (3s ramp-up, 7.5s plateau, 2.5s ramp-down), a jittered interval of 2-16 seconds, and then intensity and unpleasantness ratings, the order of which was counterbalanced. A jittered inter-trial interval (ITI) of 6-14 seconds separated the trials. gLMS = general Labeled Magnitude Scale.

**Figure S2.**
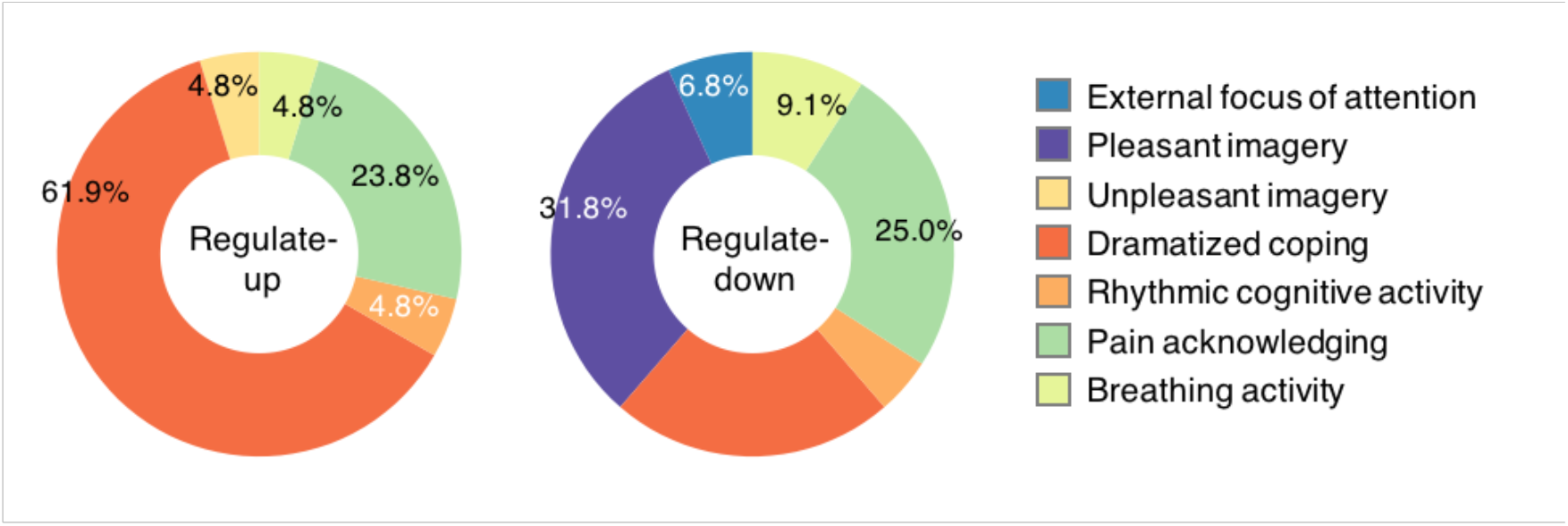
Analysis of post-experiment questionnaire on cognitive regulation strategies used by participants. The pie chart showing the proportions of regulation categories used by participants. To examine the actual regulation strategies used by participants, a post-experimental survey was administered. In the survey, we asked participants to describe the most effective strategy they used to regulate pain up or down. 38 participants responded (e.g. “For the up condition, I imagined that the thermode was burning a hole in my skin”, or “For the down, I thought of drinking a hot cup of hot chocolate on a cold day”). We then asked emotion and pain researchers (*n* = 11) to categorize these responses into one of the eight categories. Six categories were taken from Fernandez & Turk [1]: “External focus of attention”, “Neutral imagery”, “Pleasant imagery”, “Dramatized coping”, “Rhythmic cognitive activity”, “Pain acknowledging.” Plus, we added two more categories based upon the responses: “Unpleasant imagery” and “Breathing activity.” Categories are defined as the following. “External focus of attention”: strategies involving a redirection of attention away from the site of stimulation. “Neutral imagery”: strategies involving imagery of neither a pleasant nor unpleasant quality. “Pleasant imagery”: strategies centering around the use of pleasant imagery. “Unpleasant imagery”: strategies centering around the use of unpleasant imagery. “Dramatized coping”: strategies involving a dramatized reconstruction of the context in which nociception occurs. “Rhythmic cognitive activity”: strategies involving cognitive activity of a repetitive or systematized nature. “Pain acknowledging”: strategies involving a reappraisal of the nociceptive stimulation in terms of objective sensations. “Breathing activity”: regulating breathing, for example “I tried to slow down my breathing and remain calm”. We made the final decision about the regulation categories for each participant’s response based on consensus across experimenters G.M. and C.W.

**Figure S3.**
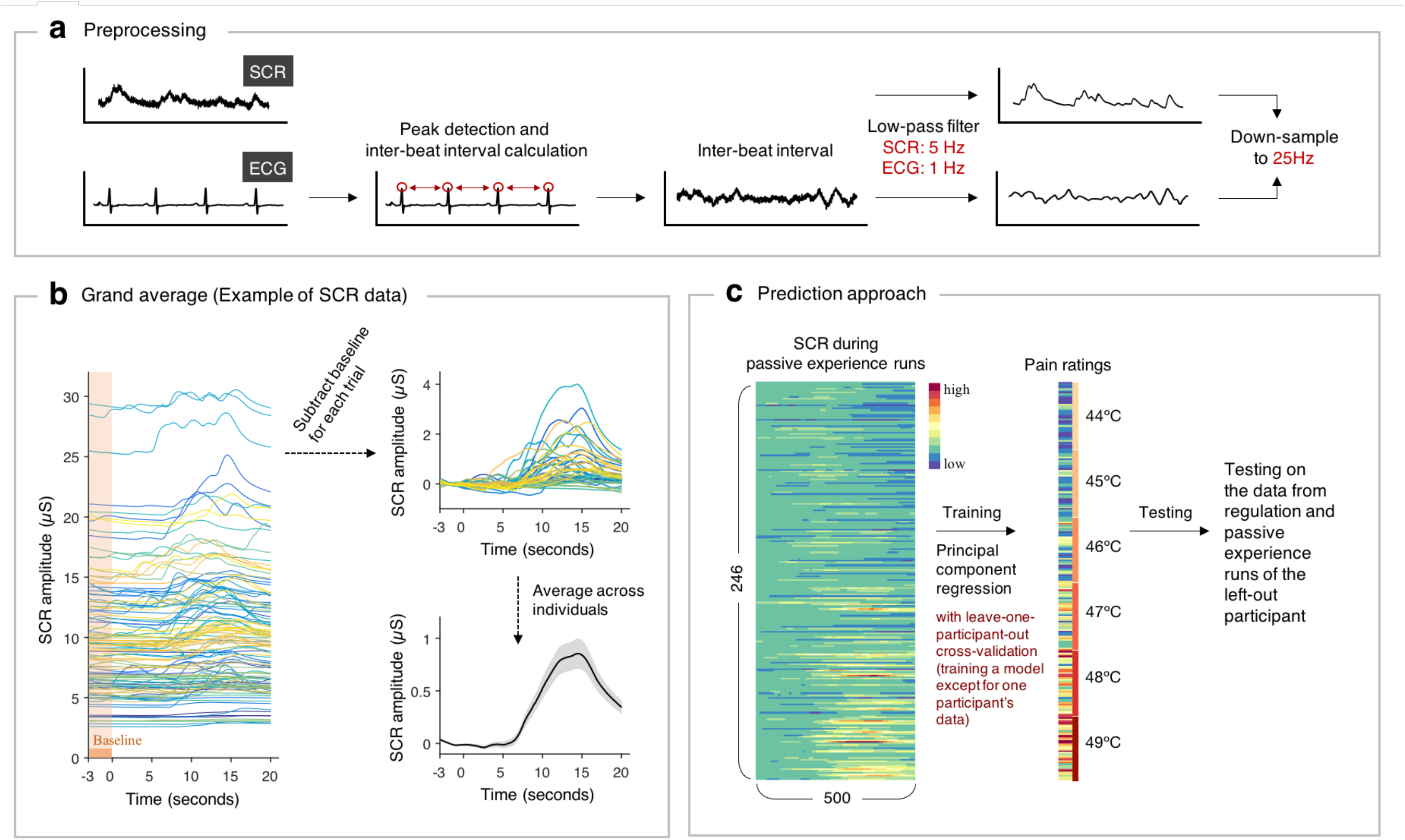
Data analysis pipeline. **(a)** Preprocessing pipeline. Electrocardiogram (ECG) data was converted into an inter-beat interval time-series. The raw SCR and IBI data were then both put through a low-pass filter (5 Hz for SCR, 1 Hz for IBI) and down-sampled to 25 Hz. **(b)** Stimulus-locked grand averages (an example of SCR data). Mean values of the three second baseline period before stimulation onset were subtracted from the stimulation epoch, and then time courses were averaged across individuals. Shades represent standard errors of the mean (s.e.m.). **(c)** Prediction approach. Using concatenated stimulus-locked average responses in only the passive experience runs as features, SCR and ECG time-course models that are predictive of pain ratings were derived with principal component regression (PCR). A leave-one-participant-out cross-validation procedure was used for testing of data from Study 1: SCR/ECG models were derived based on physiological data from passive experience conditions for all participants except for one out-of-sample participant, and the models were then tested on the out-of-sample participant’s data in all three conditions by calculating the dot-product between the time-series weights and stimulus-locked physiological data. This process was done iteratively for each participant. Note that the data from regulation runs were not included in the model developing procedure at all.

**Figure S4.**
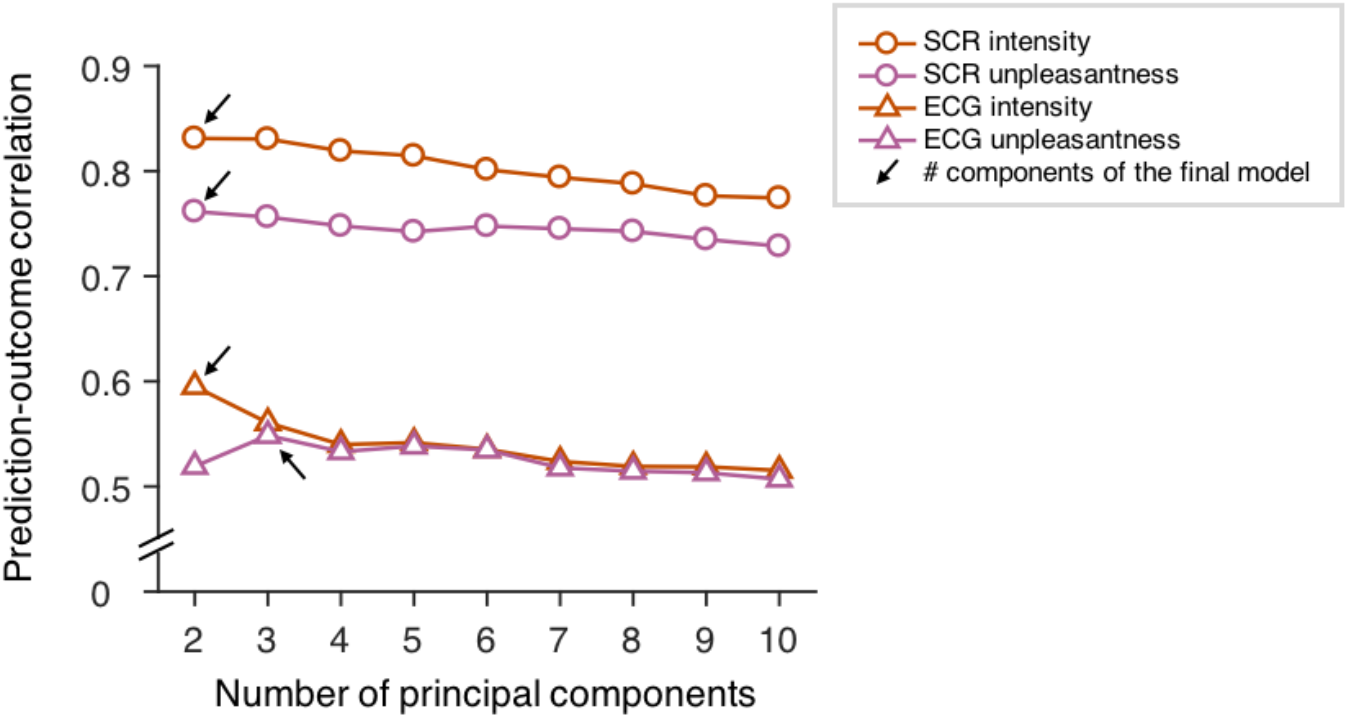
Selection of the number of components for physiological markers. Shown here are the mean prediction-outcome correlations (i.e., correlations between the actual outcome values, *y* and predicted values, 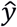) for different physiological predictive models with different numbers of components. To select the number of components for the final models that maximized the predictive performance, we used the leave-one-participant-out cross-validation procedure. The final number of components used in each model is indicated by the black arrow.

**Figure S5.**
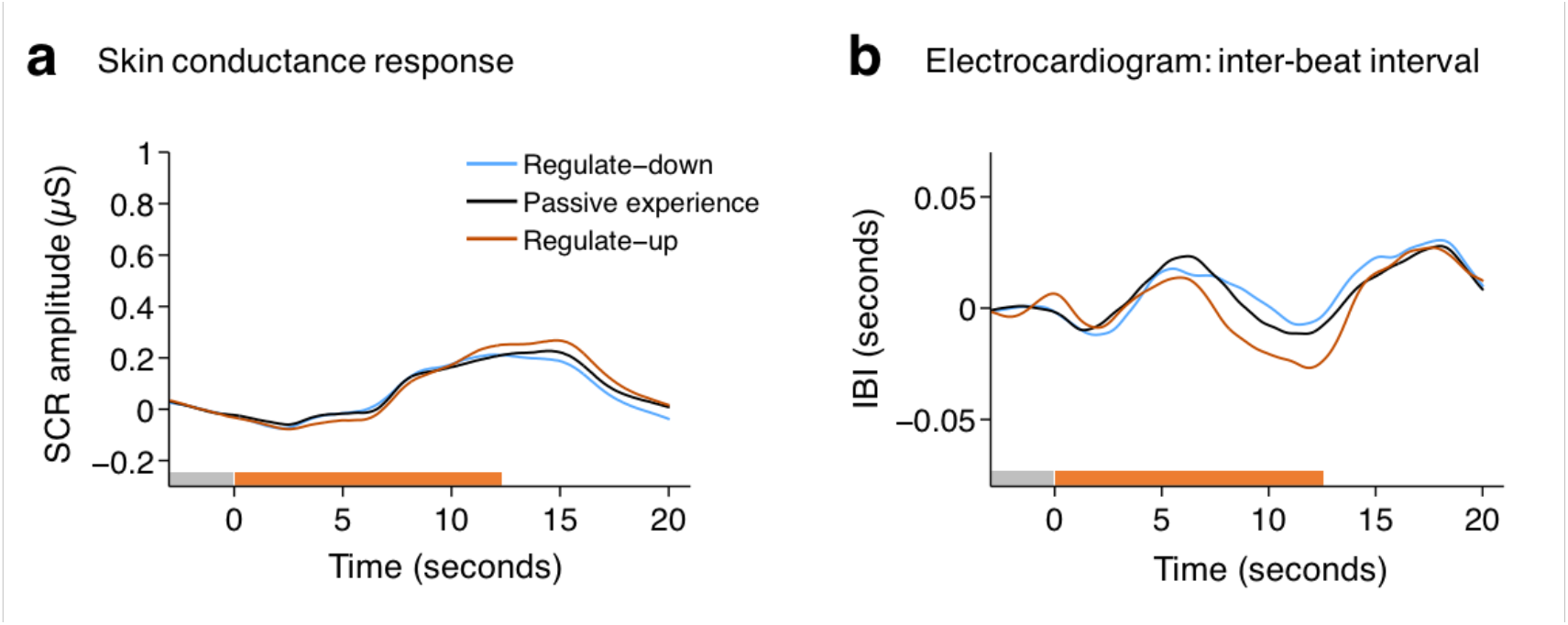
Mean physiological responses in different regulation conditions. Stimulus-locked grand average of **(a)** skin conductance responses (SCR) and **(b)** electrocardiogram’s inter-beat intervals (IBI) across participants for each regulation condition. Data from 3 seconds prior to the thermal stimulation onset were used as a baseline.

**Figure S6.**
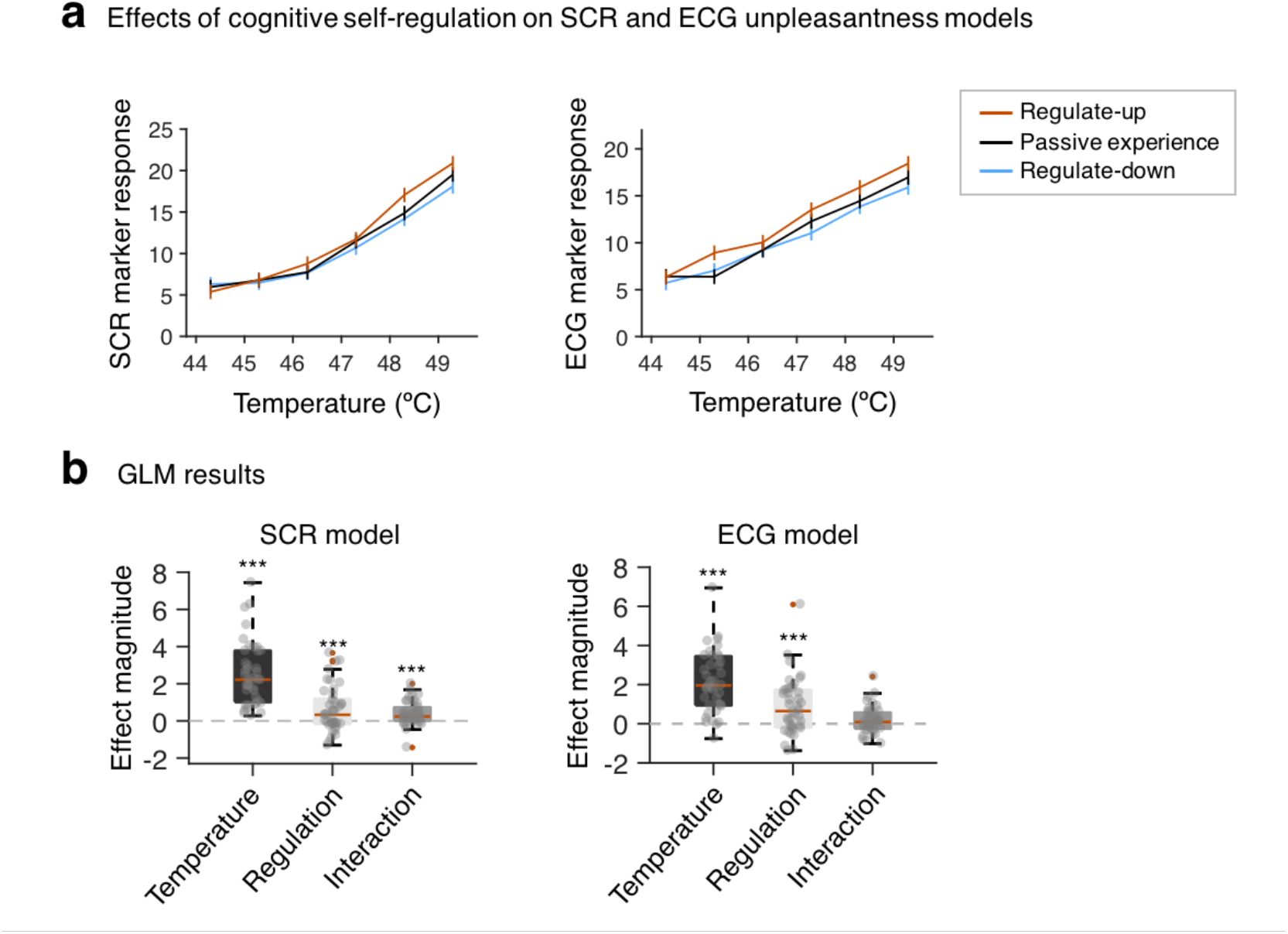
Effects of cognitive self-regulation on SCR and ECG unpleasantness markers. These are analogous plots to Fig. 5, except that these are the results for SCR and ECG unpleasantness markers. **(a)** Predicted pain scores by SCR and ECG pain unpleasantness models. Error bars represent within-subject S.E.M. **(b)** Multi-level general linear model results. Both stimulus intensity and cognitive self-regulation had significant effects on SCR and ECG unpleasantness marker responses. \*\*\**p* < .001; Bootstrap tests (10,000 iterations) were used for significance testing.

**Figure S7.**
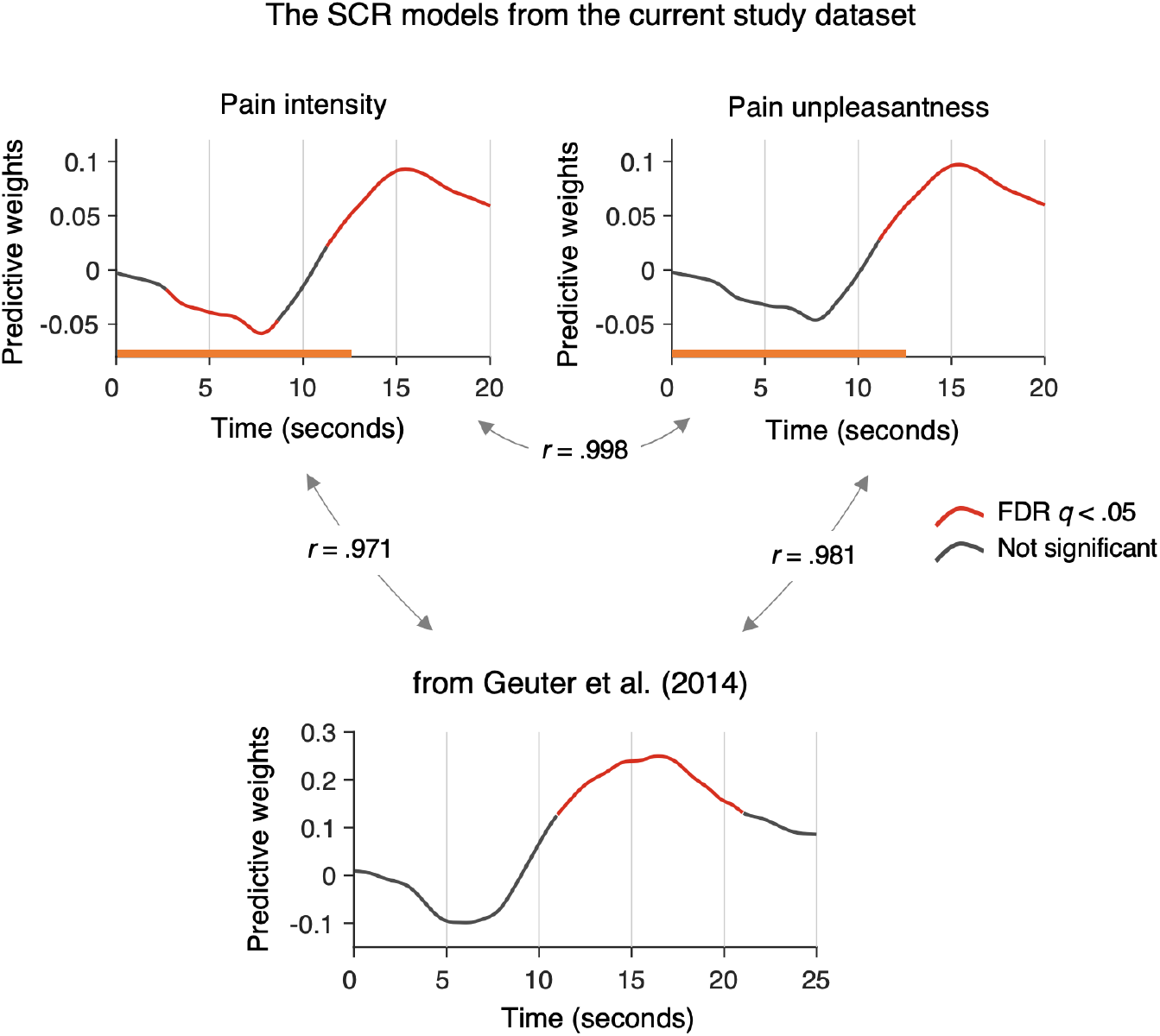
Testing the SCR model from Geuter et al. [2]. In these analyses, we used the SCR model based on 17-seconds heat stimuli.

1. We first compared Geuter’s SCR model to the models form the current study. Despite the differences in stimulus duration, Geuter’s SCR model was highly similar to our SCR models in terms of peak locations and significant time points.
2. We also tried to predict the pain ratings with the Geuter’s SCR model. The predictive performance was comparable to our models; the mean prediction-outcome correlations for the passive experience runs were *r* = .82 ± 0.027, *p* < .0001 for intensity ratings and *r* = 0.67 ± 0.060, *p* < .0001 for unpleasantness ratings. For regulation runs, the mean prediction-outcome correlations were *r* = .81 ± 0.022, *p* < .0001 for intensity ratings and *r* = 0.71 ± 0.041, *p* < .0001 for unpleasantness ratings.
3. We lastly tested whether cognitive self-regulation has a significant effect on Geuter’s SCR model. Similar to our main results, both stimulus intensity and self-regulation had significant effects on Geuter’s SCR model response, 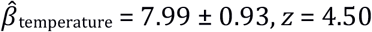, *z* = 4.50, *p* < .0001, 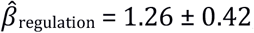, *z* = 3.44, *p* < .001.

**Table S1.** Multi-level general linear model results for different contrasts.

| Outcome (Y) | Contrasts (X) | *beta* | s.e.m | bootstrap test results |  |
| --- | --- | --- | --- | --- | --- |
|  |  |  |  | *z* | *p* |
| pain intensity ratings | stimulus intensity (temperature)<sup>a</sup> | 5.01 | 0.32 | 3.70 | 0.0001 |
|  | regulate-up vs. regulate-down<sup>a</sup> | 2.12 | 0.36 | 3.75 | 0.0001 |
|  | regulate-up vs. passive experience<sup>b</sup> | 1.44 | 0.52 | 2.46 | 0.0069 |
|  | passive experience vs. regulate-down<sup>c</sup> | 2.80 | 0.56 | 4.02 | <.0001 |
| pain unpleasantness ratings | stimulus intensity (temperature)<sup>a</sup> | 5.50 | 0.38 | 3.49 | 0.0002 |
|  | regulate-up vs. regulate-down<sup>a</sup> | 5.19 | 0.68 | 4.44 | <.0001 |
|  | regulate-up vs. passive experience<sup>b</sup> | 4.77 | 0.77 | 3.80 | 0.0001 |
|  | passive experience vs. regulate-down<sup>c</sup> | 5.62 | 1.02 | 5.50 | <.0001 |
| SCR intensity marker | stimulus intensity (temperature)<sup>a</sup> | 2.50 | 0.33 | 4.76 | <.0001 |
|  | regulate-up vs. regulate-down<sup>a</sup> | 0.61 | 0.18 | 4.12 | <.0001 |
|  | regulate-up vs. passive experience<sup>b</sup> | 0.70 | 0.38 | 1.64 | 0.0500 |
|  | passive experience vs. regulate-down<sup>c</sup> | 0.51 | 0.30 | 1.53 | 0.0630 |
| SCR unpleasantness marker | stimulus intensity (temperature)<sup>a</sup> | 2.78 | 0.38 | 4.91 | <.0001 |
|  | regulate-up vs. regulate-down<sup>a</sup> | 0.61 | 0.19 | 3.95 | <.0001 |
|  | regulate-up vs. passive experience<sup>b</sup> | 0.72 | 0.42 | 1.55 | 0.0604 |
|  | passive experience vs. regulate-down<sup>c</sup> | 0.49 | 0.33 | 1.19 | 0.1162 |
| ECG intensity marker | stimulus intensity (temperature)<sup>a</sup> | 1.50 | 0.23 | 4.85 | <.0001 |
|  | regulate-up vs. regulate-down<sup>a</sup> | 0.62 | 0.15 | 3.91 | <.0001 |
|  | regulate-up vs. passive experience<sup>b</sup> | 1.09 | 0.31 | 3.38 | 0.0004 |
|  | passive experience vs. regulate-down<sup>c</sup> | 0.14 | 0.30 | 0.52 | 0.6968 |
| ECG unpleasantness marker | stimulus intensity (temperature)<sup>a</sup> | 2.27 | 0.29 | 4.13 | <.0001 |
|  | regulate-up vs. regulate-down<sup>a</sup> | 0.87 | 0.23 | 4.09 | <.0001 |
|  | regulate-up vs. passive experience<sup>b</sup> | 1.24 | 0.43 | 2.82 | 0.0024 |
|  | passive experience vs. regulate-down<sup>c</sup> | 0.50 | 0.45 | 0.58 | 0.2800 |

*Note.* This table shows results from three different multi-level general linear models (MGLM) represented by different superscripts. All three MGLM models include three types of independent variables: (i) stimulus intensity (temperature, °C), (ii) regulation, and (iii) the interaction term between the stimulus intensity and regulation. For three different MGLM models, we used different contrasts for the “regulation” variable. <sup>a</sup> For the regulate-up vs. regulate-down contrast, the regulation was coded as 1, 0, and -1 for regulate-up, passive experience, and regulate-down, respectively. <sup>b</sup> For the regulate-up vs. passive experience contrast, the regulation regressor was coded as 0.5 and -0.5 for regulate-up and
passive experience, respectively to make the unit consistent with the other models. <sup>c</sup> For the passive experience vs. regulate-down contrast, the regulation regressor was coded as 0.5 and -0.5 for passive experience and regulate-down. For model <sup>b</sup> and <sup>c</sup>, we only report the results of the regulation contrasts in this table.

